# Targeted Antisense Oligonucleotide Treatment Rescues Developmental Alterations in Spinal Muscular Atrophy Organoids

**DOI:** 10.1101/2025.01.17.633436

**Authors:** Irene Faravelli, Paola Rinchetti, Monica Tambalo, Illia Simutin, Lisa Mapelli, Sara Mancinelli, Matteo Miotto, Mafalda Rizzuti, Andrea D’Angelo, Chiara Cordiglieri, Giulia Forotti, Clelia Peano, Paolo Kunderfranco, Luca Calandriello, Giacomo P. Comi, Elvezia Paraboschi, Eleonora Pali, Francesca Beatrice, Egidio D’Angelo, Serge Przedborski, Monica Nizzardo, Simona Lodato, Stefania Corti

## Abstract

Spinal muscular atrophy (SMA) is a severe neurological disease caused by mutations in the *SMN1* gene, characterized by early onset and degeneration of lower motor neurons. Understanding early neurodevelopmental defects in SMA is crucial for optimizing therapeutic interventions. Using spinal cord and cerebral organoids generated from multiple SMA type I donors, we revealed widespread disease mechanisms beyond motor neuron degeneration. Single-cell transcriptomics uncovered pervasive alterations across neural populations, from progenitors to neurons, demonstrating SMN-dependent dysregulation of neuronal differentiation programs. Multi-electrode array analysis identified consistent hyperexcitability in both spinal and brain organoids, establishing altered electrical properties as a central nervous system-wide feature of pathogenesis. Early administration of an optimized antisense oligonucleotide (ASO) that restored SMN levels rescued morphological and functional deficits in spinal cord organoids across different genetic backgrounds. Importantly, this early intervention precisely corrected aberrant splicing in newly identified *SMN1* targets enriched at critical nodes of neuronal differentiation. Our findings demonstrate that early developmental defects are core features of SMA pathogenesis that can be prevented by timely therapeutic intervention, providing new insights for optimizing treatment strategies.

## Introduction

Spinal muscular atrophy (SMA) is a severe inherited neurological disorder and still one of the leading genetic causes of death in childhood^1,2^. SMA is caused by autosomal recessive mutations in *SMN1*, resulting in reduced expression of full-length SMN protein^3^. This leads to progressive degeneration of more vulnerable spinal motor neurons (MNs), causing proximal muscle weakness and atrophy, and early life-threatening respiratory failure^4,5^. SMA phenotypic spectrum is very broad and consists of different clinical forms, from fatal to mild, which are ranked on the base of the motor milestones reached by the patients^6^.

The SMN protein is expressed in several fetal tissues including the brain, spinal cord, skeletal muscle, heart, and kidney; it significantly decreases after birth, suggesting that SMN is particularly important during prenatal life^7,8^. Recent data from human SMA autopsies associated abnormal MN axon development *in utero* with postnatal degeneration in SMA^9^. Beyond spinal cord involvement, defects in cognitive function and brain neuropathology have been reported in SMA patients, but only partially documented^10^. The early events that drive the degeneration as well as their impact on the entire central nervous system (CNS) are still largely unknown: considering pre-symptomatic factors is key when modeling the disease.

Recently, FDA/EMA approved breakthrough therapeutic strategies to restore the levels of SMN and they are currently in clinical use^5,11^. These approaches are based on either cDNA replacement of *SMN1* or the use of antisense oligonucleotides (ASOs) or small molecules to achieve splicing correction of *SMN2*. *SMN2* is a paralog of *SMN1* that produces a truncated and rapidly degraded SMN protein, which can be used as point of action to restore SMN levels^12^. These drugs have dramatically changed the clinical prognosis of SMA, and for the fatal to severe SMA type 1, are able to reduce the disease to a chronic condition by halting or preventing motor disability. Although growing clinical data show that approved treatments also provide benefits in the chronic phase, it is emerging that the greatest impact is achieved when patients are treated pre-symptomatically^5^. Remaining unmet therapeutic needs are the treatment of patients after symptom onset^5^ and the elusive understanding of the factors halting treatment efficacy. Critically, even newborns identified through screening show incomplete motor recovery, particularly those with two *SMN2* copies^13^. Therefore, a better understanding of SMA mechanisms in prenatal disease stages is pivotal to optimizing current therapeutic approaches with the aim of increasing efficiency, targeting biodistribution, and developing new clinical strategies.

Conventional 2D cultures and animal models can only partially recapitulate the phenotypic and genetic complexity of human MN diseases. CNS 3D human organoids^14–16^, in contrast, display a high degree of cellular diversity, enabling them to model complex cell–cell interactions, while modeling the patient-specific genetic background.

Here, we leveraged a novel and efficient protocol for the generation of induced pluripotent stem cell (iPSC)-derived spinal cord organoids (SCOs) and unveiled early clinically relevant SMA pathological phenotypes. We detected alterations in progenitors’ behaviour, early neuronal differentiation defects, even before hallmarks of cell death appeared, and identified disease susceptibility for distinct neuronal subtypes, beyond MNs.

A pervasive alteration of the splicing machinery has been involved in several neurodegenerative diseases, and it has been recently postulated for the adult MN disease amyotrophic lateral sclerosis (ALS)^17^. SMN protein has a recognized role in the splicing machinery, but the impact of its disruption on developmental processes is still unexplored. Using SCOs, we identified genes with abnormal splicing in SMA and those rescued by SMN-targeting therapies. This allowed us to study how restoring SMN reshapes the transcriptional landscape in patient-derived SCOs. Interestingly, we found that SMN deficiency disrupts the splicing of key genes involved in neural differentiation, potentially triggering SMA pathogenesis. Our findings demonstrate SMN critical role in neural development and the significance of its restoration in therapeutic strategies.

## Results

### SMN Deficiency Impacts Progenitors and Neurons in the Spinal Cord

We derived human SCOs from healthy control (CTRL, *n* = 5) and SMA type 1 patient (*n* = 5) induced pluripotent stem cell (iPSC) lines. Spinal cord development is a complex process that entails both rostro-caudal specification, mediated by Wnt and retinoic acid (RA), and dorso-ventral patterning, downstream of sonic hedgehog (SHH) signaling^18,19^. Thus, SCOs were cultured in the presence of small molecules acting on these pathways to recapitulate *in vitro* the spinal cord specification^20–22^ (**Fig. 1A**). In both CTRL and SMA organoids, the treatment promoted a progressive caudal neuralization with the appearance at the early differentiation stage (1 month) of neural progenitors (**Fig. 1B**) expressing specific markers including PAX6 and NESTIN, followed by the appearance of the neuronal markers DCX and MAP2 (**Fig. 1B**). As the differentiation proceeded, we observed the presence of spinal precursors expressing HOXB4 and OLIG2 along with ISL1- and MNX1-positive MNs (**Fig. 1B**). Together, the data show that SCOs can recapitulate key aspects of spinal cord neurogenesis.

**Figure 1.**
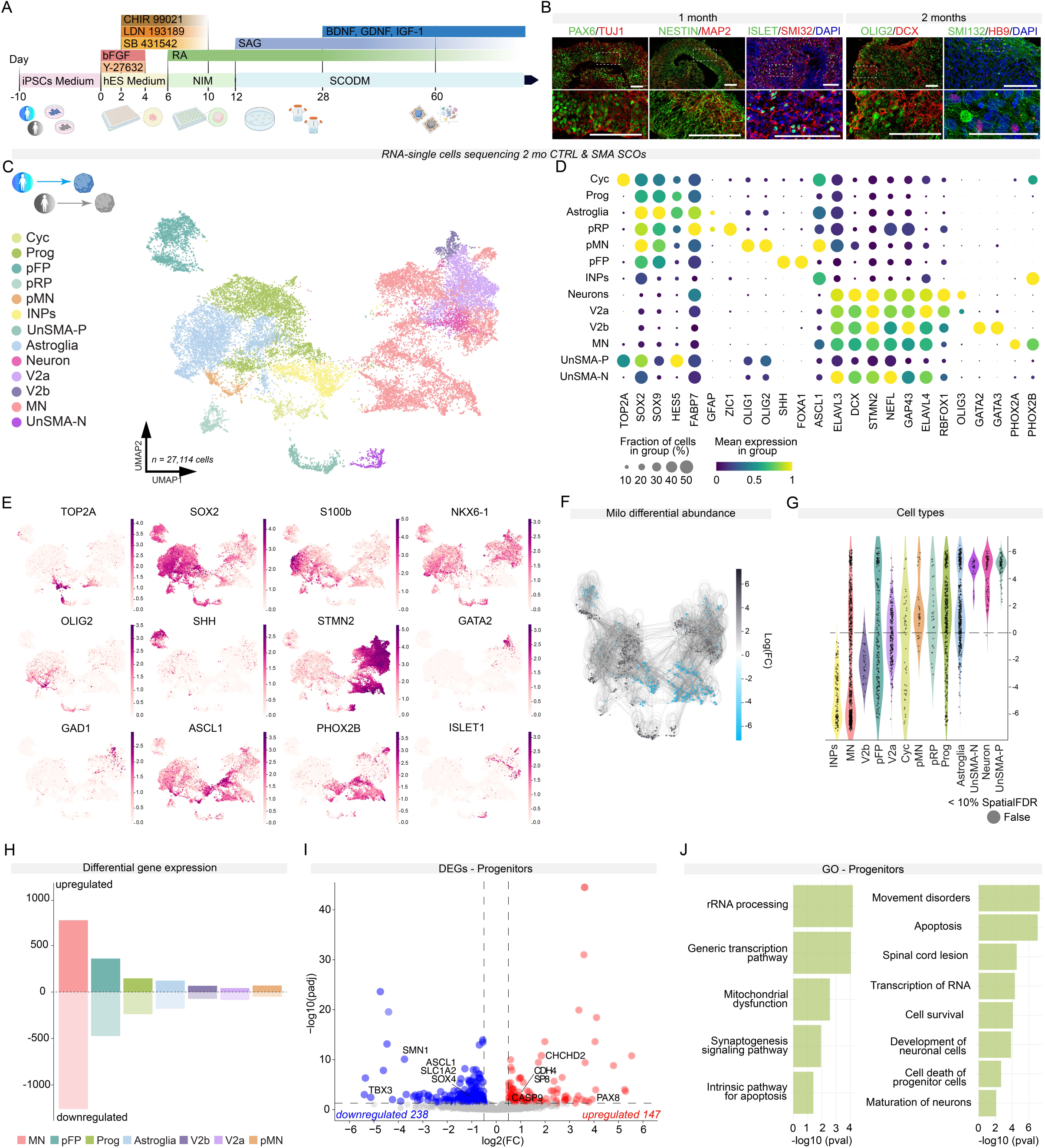
CTRL and SMA SCOs display different subtypes of spinal cord cell populations. **A**, Schematics of human SCO generation. **B**, Immunostaining of developmental and neuronal markers in SCOs at 1 and 2 months. SCOs exhibit positive staining for neurodevelopmental markers, including PAX6, NESTIN, and OLIG2, as well as neuronal markers such as TUJ1, MAP2, SMI132, and DCX and motor neuron markers such as ISLET1 and HB9. Nuclei were stained with DAPI (blue). Scale bar: 100 µm. **C**, CTRL and SMA single cell RNA-seq at 2 months (mo). Unsupervised UMAP subdivides SCOs into neural progenitors (cycling cells (Cyc), progenitors (Prog), progenitors roof plate (pRP) intermediate neuronal progenitors (INPs)), floor plate progenitors (pFP), motor neuron progenitors (pMN) and motor neurons (MN), differentiated neuronal cells (Neuron), subtypes of ventral neurons (V2a, V2b), astroglia, and a cluster of unmatched SMA cells (unmatched-SMA progenitors (UnSMA-P), unmatched-SMA neurons (UnSMA-N). **D**, Dot plots showing cluster-specific expression of representative genes associated with cell differentiation; dot size corresponds to the percentage of cells in a cluster expressing each gene and the color corresponds to the average expression level (from violet to yellow, low to high). **E**, UMAP feature plots showing the log-normalized counts of selected representative genes for proliferative markers (*TOP2A* and *SOX2*), progenitor markers (*NKX6-1, OLIG2*, *SHH*, *GATA2*) and neuronal differentiation markers (*DCX*, *STMN2*, *PHOX2B* and *ISL1*). **F**, Abstracted graph of neighborhoods of the results from Milo differential abundance testing. SMA enrichment in grey and CTRL enrichment in blue. **G**, Violin plot showing the distribution of log-fold changes of differentially abundant neighborhoods assigned to cell type labels. **H**, Numbers of deregulated genes across different clusters. **I**, Volcano plot of differential gene expression in progenitor clusters in SMA SCOs relative to CTRL. Values of log2 fold change are plotted against -log10 of the q-value for each gene. Red denotes genes upregulated in SMA while blue denotes genes downregulated in SMA (absolute log fold change ≥0.2 min.pct = 0.25 and adjusted q-value< 0.05, thresholds plotted as dashed lines). **J**, Selected Gene Ontology for upregulated and downregulated genes in progenitor cluster performed with QIAGEN Ingenuity Pathway Analysis (IPA) software.

To explore the early events driving SMA pathogenesis, we analyzed 2-month-old CTRL and SMA SCOs composition by single-cell RNA-seq (**Fig. 1C-E** and **Suppl. Fig. 1**). Accounting for technical variability, we sequenced individual SCOs deriving from 2 CTRL and 2 SMA lines and successfully profiled a total of 27,114 cells. Module annotation with defined sets of marker genes for dorsal and ventral domains identified different classes of neural progenitors (Prog, *SOX2, SOX9*), including floor plate progenitors (pFP, *SHH*), differentiating neurons (Neuron, *DCX*), and subtypes of spinal cord neuronal domains (**Fig. 1C-E** and **Suppl. Fig. 1**). At this stage, we could also identify distinct clusters of MN precursors (pMN, *OLIG2*), intermediate precursors (INPs, *ASCL1*) and postmitotic MNs (MN, *PHOX2B, ISLET1*) whose transcriptional profiles could be integrated with early endogenous datasets^23^ (**Fig. 1C-E** and **Suppl. Fig. 1C**). The selective expression of defined combinations of HOX transcripts corresponding to a posterior neural tube identity suggested the acquisition of a cervical spinal cord signature (**Suppl. Fig. 1D)**. Overall, these results confirm that SCOs represent robust tools for modeling neurological diseases with multicellular involvement, such as SMA.

To investigate whether cell composition and relative contribution of neuronal types are altered in the absence of SMN protein, we performed a differential abundance test using the MiloPy framework^24^ (**Fig. 1F-G**). This revealed that specific cell subtypes were differentially represented in SMA organoids, with MNs significantly decreased in SMA genotype. Interestingly, in contrast with the reduction of mature MNs, pMNs, were enriched in SMA organoids, as well as other clusters of neural progenitors (**Fig. 1F-G)**. These results support a broader effect on neural diversity in the absence of SMN, which impacts the proportion of different cell types in the developing organoids.

We, therefore, investigated the transcriptional changes associated with the lack of SMN in each cluster. To account for the differences in abundance presented in SMA SCOs, we performed differential gene expression analysis in clusters presenting at least 25% of cells of each genotype (**Fig. 1H** and **Suppl. Fig. 2;** absolute log fold change ≥ 0.5 and adjusted p-value < 0.05). While the largest proportion of deregulated genes (2031 genes differentially expressed between CTRL and SMA SCOs, 772 upregulated and 1259 downregulated) was observed in cells ascribed to the MN cluster, progenitor cells were the next most affected. Interestingly, specific genes related to progenitor fitness were downregulated (*ASCL1*, *SOX4)*, while *CASP9,* involved in the apoptosis signaling, was upregulated (**Fig. 1I**). Gene ontology analysis on progenitors (**Fig. 1J**) revealed a pervasive effect of the lack of SMN on pathways involved in several developmental processes. Overall, these findings highlight the profound impact of SMN deficiency on the cellular composition and transcriptional landscape of SCOs. While the significant reduction in spinal MNs is a key hallmark of SMA, the altered abundance and transcriptional dysregulation of progenitor cells reveal an equally critical aspect of the disease pathology with a broader disruption in neural development, often disregarded in the investigation of this disease.

### SMA progenitor defects precede motor neuron impairment

To longitudinally capture the differentiation progression phenotype at the cellular level, we performed high-content image analysis (**Fig. 2A-D**). Notably, aberrant differentiation trajectories were observed in SMA SCOs. While CTRL SCOs showed a clear reduction in SOX2-expressing neural progenitors and an increase in the post-mitotic neuronal marker TUJ1 from 1 to 2 months, SMA SCOs exhibited a less pronounced reduction in SOX2 and no increase in the proportion of TUJ1-expressing cells (**Fig. 2A-D**; percentage of SOX2 positive cells in each organoid at 1 *vs.* 2 months: CTRL 47.57% ± 11.94 *vs.* 20.40% ± 13.85, p<0.0001; SMA 38.79% ± 14.42 *vs.* 25.44% ± 13.91, p=0.0148; percentage of TUJ1 positive cells in each organoid at 1 *vs.* 2 months: CTRL 13.52% ± 4.788 *vs.* 28.42% ± 12.35, p=0.0002; SMA 21.49% ± 14.28 *vs.* 13.84% ± 6.759, p=0.0764). This suggests an early defect in differentiation and maturation.

**Figure 2.**
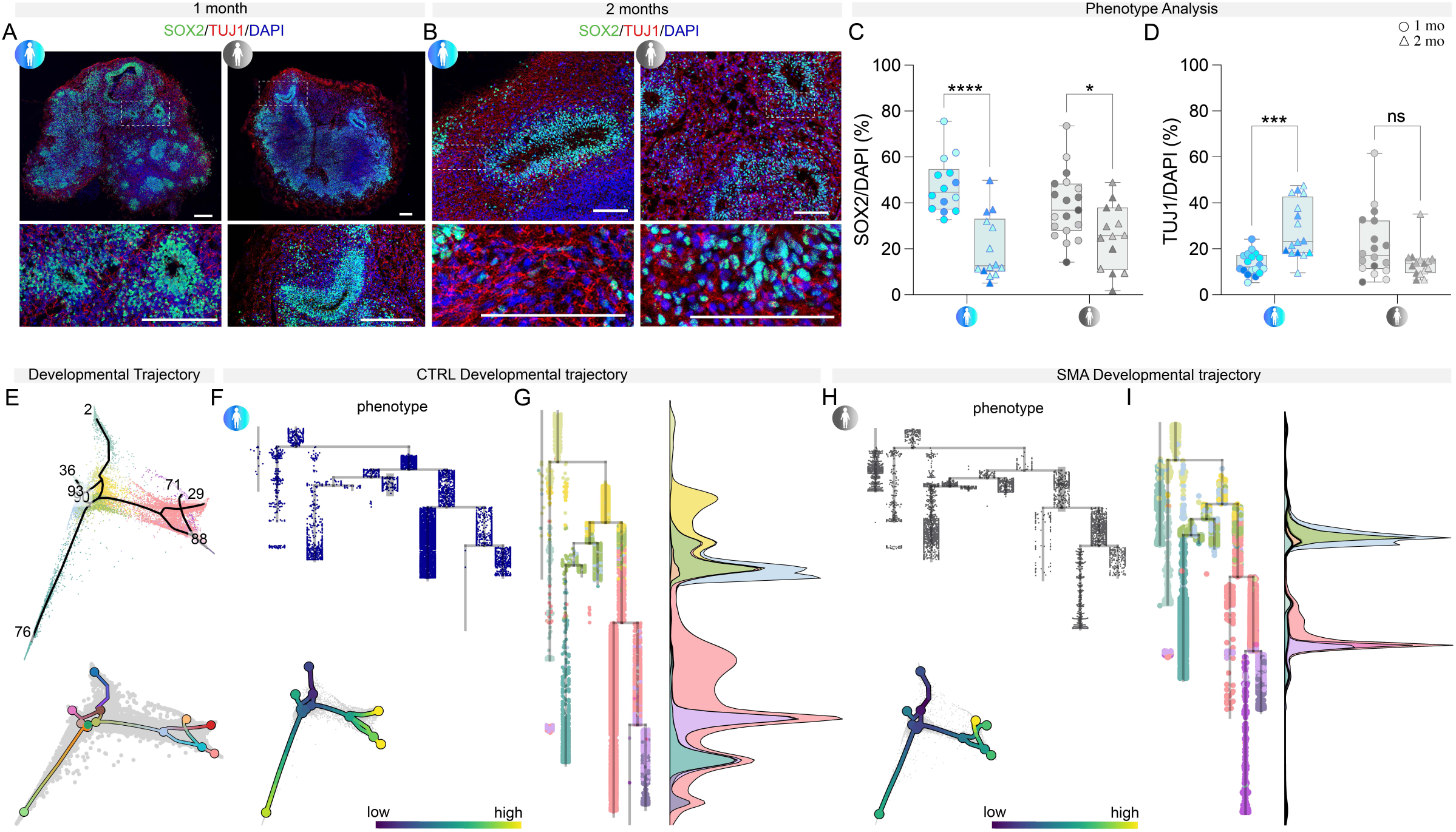
Neurodevelopmental disease signatures of SMA SCOs. **A-B**, Representative images of SOX2 (green) and TUJ1 (red) staining at 1 and 2 mo in CTRL and SMA SCOs. Scale bar: 200 µm. Nuclei were stained with DAPI (blue). **C**, Quantification of SOX2-positive cells (as a percentage of cells revealed by DAPI staining) at 1 mo (circles) and 2 mo (triangles) (two-way ANOVA, Sidak’s multiple comparisons test: *p*<0.0001 and *p*=0.0148 for CTRL and SMA, respectively). **D**, Expression of TUJ1 increase over time in the CTRL, meanwhile in the SMA SCOS there is a trend of decreasing in the expression of TUJ1 (two-way ANOVA, Sidak’s multiple comparisons test: *p*=0.0002 and *p*=0.0764 for CTRL and SMA, respectively). **E**, Principal graph inferred on Palantir’s diffusion components with SimplePPT^25^ for both genotypes. **F-I**, CTRL and SMA SCOs developmental trajectory. Pseudotime was calculated using cycling cells as the root, and the common trajectory was subsequently separated for CTRL and SMA SCOs. Cells are colored according to the pseudotime and cell-type labels and visualized in a form of the dendrogram and pseudotime density plot.

Since SCOs recapitulate the early stage of spinal cord development, and our data suggest that the physiological process of cell death could be enhanced in SMA, we longitudinally tested CTRL and SMA SCOs for programmed cell death. SMA SCOs indeed exhibited a difference in apoptosis dynamics in comparison with CTRL SCOs, with a steep enhancement at 2 months (percentage of apoptotic nuclei in each organoid at 1 *vs*. 2 months: CTRL 11.30% ± 2.512 *vs.* 7.369% ± 5.125, *p* = 0.3786; SMA, 4.245% ± 2.372 *vs.* 23.02% ± 11.34, *p* < 0.0001; **Suppl. Fig. 3A-B**). Enrichment analysis of apoptosis-related biological functions, based on differentially expressed genes between SMA and CTRL SCOs, revealed that these processes were predominantly upregulated in SMA pMNs and MNs (**Suppl. Fig. 3C**).

Differences in cell death rate might explain, at least in part, the reduced MN population in SMA SCOs captured by MiloPy (**Fig. 1H**), while the altered composition with enriched progenitors and neurons suggests changes in early developmental programs in SMA tissue.

To capture the differences in developmental dynamics from progenitors to neurons, we calculated the developmental trajectory using sc_Fates package^25^ (**Fig. 2E-I**). Cycling populations were identified as roots for tree inference for both conditions and distinct tree topologies were observed. For the CTRL organoids, the pseudo-temporal ordering of cell types approximated that of human fetal development^26^, following a differentiation trajectory that progressed from progenitors to intermediate precursors, neuronal cell types and terminally commitment of MNs (**Fig. 2F-G**). The cells split into two major branches, one containing progenitors in distinct states of their proliferating activity and committed progenitors of the floor plate, and one containing neural progenitors toward differentiated neurons (**Fig. 2F-G**). SMA SCOs showed a strikingly different distribution of cell states, with increased distribution of cells toward the starting and early segments of the developmental trajectory, i.e., at an intermediate stage of development. This was accompanied by a significant reduction in the endpoint of the trajectory, represented by maturing neuronal subtypes (**Fig. 2H-I**).

Altogether, these data support our previous results pointing at an early alteration in SMA neurodevelopment, where cells pause at intermediate developmental states unable to proceed along their maturation, which could, in turn, impact in their ability to survive in the nascent circuits.

### SMA motor neuron developmental trajectory is hindered

To investigate gene expression changes during MN differentiation, we partitioned the dataset to focus specifically on MN linage, including pMNs, INPs, and MNs (**Fig.3** and **Suppl. Fig. 4**). *De novo* clustering of the MN lineage partition revealed eight distinct clusters (**Suppl. Fig. 4A**), with MN cells being more abundant in CTRL genotypes (**Suppl. Fig. 4B**). Within the MN clusters, we could further distinguish five individual clusters based on the expression levels of *PHOX2B* versus *ISL1/MNX1* (**Suppl. Fig. 4C**).

**Figure 3.**
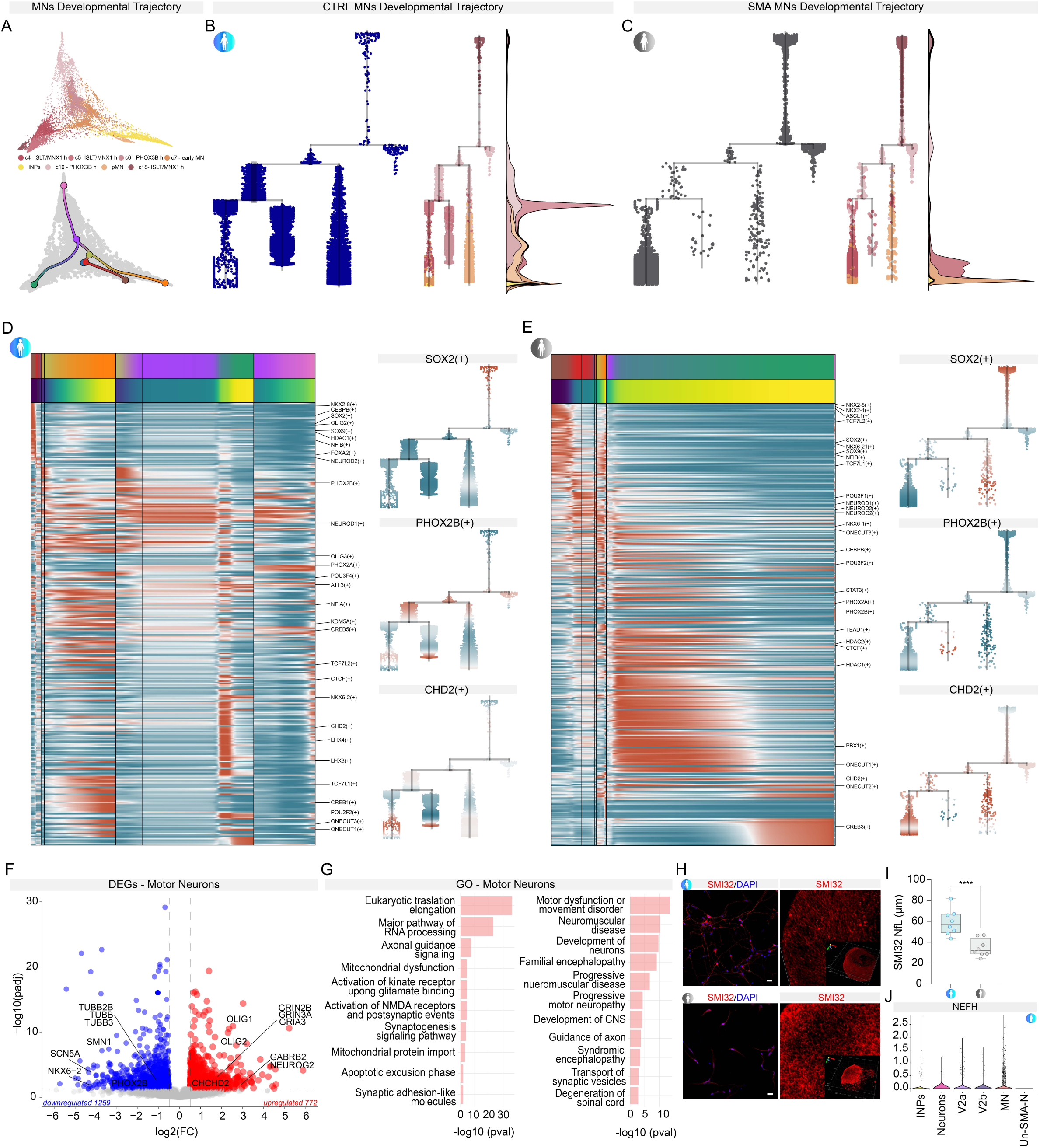
SMA SCOs show molecular signatures of disease. **A**, Common motor neurons (MNs) developmental trajectory for CTRL and SMA SCOs projected on the multiscale diffusion space. **B**-**C**, The trajectories of CTRL and SMA SCOs motor neurons were separated, as shown in the dendrogram and pseudotime distribution. **D-E**, Heatmap displaying regulon activity trends and pseudotime associated with the CTRL (**D**) and SMA (**E**) trajectories, binned by each trajectory segment. Regulon activity scores of *SOX2*, *PHOX2B*, and *CHD2* in CTRL and SMA samples. High level of regulon’s activity in red and low level of in blue. **F**, Volcano plot of differential gene expression in MN clusters in SMA SCOs relative to CTRL. Values of log2 fold change are plotted against -log10 of the q-value for each gene. Red denotes genes upregulated in SMA while blue denotes genes downregulated in SMA (absolute log fold change ≥0.2 min.pct = 0.25 and adjusted q-value <0.05, thresholds plotted as dashed lines). **G**, Selected Gene Ontology for upregulated and downregulated genes in MNs performed with QIAGEN Ingenuity Pathway Analysis (IPA) software. **H**, Dissociated SMA organoids had a reduced SMI32-stained neurite length in culture. Scale bar: 20 µm. Nuclei were stained with DAPI (blue). The panel in the bottom left corner represents the whole-mount followed by two-photon acquisition of SMI32 staining of CTRL and SMA SCOs. **I**, Quantification of (**H**). Neurite length is significantly lower in SMA than CTRL SCOs (unpaired t-test, *p*<0.0001). **J**, Violin plot showing the enriched cells for neurofilament heavy chain (*NEFH*).

Next, we explored developmental pathways within the MN partition, allowing us to map the neurogenic progression from pMN to MNs (**Fig. 3A-C**). In CTRL SCOs, this neurogenic trajectory followed the expected sequential progression of gene expression during differentiation (**Fig. 3B**). Progenitor genes, including *OLIG2*, were expressed earlier in pseudotime, followed by proneural genes such as *NEUROD1*, and finally, the postmitotic neuron marker *PHOX2B* was detected at the latest pseudotime points (**Fig. 3A, B**). A different scenario emerged in the MN differentiation trajectory of SMA SCOs. In contrast to the continuous progression seen in CTRL samples, the differentiation trajectory in SMA appeared polarized. This polarization was characterized by a concentration of cells at the progenitor state at the beginning of the trajectory, an accumulation of cells in an intermediate state, and a notable enrichment of MNs at the end of the trajectory, predominantly within a single branch **(Fig. 3C)**. This suggests that SMN deficiency may disrupt the progressive acquisition of fate along MN differentiation, leading to an imbalance between progenitors and terminally defined neurons.

The interplay between specific transcription factors (TFs) is pivotal for the execution of correct developmental programs in the spinal cord^27^; cell-type-specific TFs represent central hubs in gene regulatory networks and alteration of their expression is associated with downstream defects in cell identity acquisition and maintenance^18^. To investigate whether the early developmental defects associated with SMA disease were driven by changes in TF activity, we used pySCENIC (Single-Cell Regulatory Network Inference and Clustering) algorithm^28^ as a tool for the selection of key TFs along the developmental trajectory of CTRL and SMA SCOs organoids (**Fig. 3D, E**). Interestingly, we found several TFs known to control key aspects of the molecular development of the human spinal cord (i.e. *SOX2, SOX9, OLIG2, NKX6-1)*) expressed along the different states of the trajectories in SMA and CTRL SCO, suggesting important variations in the activity of transcriptional programs among cell states in absence of SMN. Histone modifiers and regulators of chromatin architecture were also identified (*CHD2, HDAC2, KDM5A*), which could be novel important determinants in human spinal cord development and could impact SMA disease course.

To determine whether the developmental TF expression implied differential gene regulatory programs, we calculated regulon activity. We observed that shared regulons of TFs key for MN differentiation (e.g., *SOX2, PHOX2B, CHD2*) had shifted profile of activity in SMA, with a sustained activity of *SOX2*-transcriptional programs, which could be associated with the prolonged maintenance of progenitor states (**Fig. 3D, E**). Indeed, dimensionality reduction plots based on SCENIC regulons activity confirmed a blend distribution of the cellular states of SMA MNs lineage, supporting the idea of altered boundaries between cell fate transitions (**Fig. 3D, E**). Together these analyses support the idea that, not only SMA SCOs fail to properly differentiate into mature MNs, but also show that the neuronal developmental trajectory, with the associated transcriptional regulations, is altered from the early progenitor state. This might impinge on MN survival as well as influence the overall spinal cord neuronal differentiation.

We then looked at the transcriptional landscape of SMA vs CTRL MNs (**Fig. 3F, G**). SMA MNs showed downregulation of genes involved in neuronal growth and synaptic maturation *(TUBB3, TUBB, SCN5A)*. Notably, a small subset of genes showed consistent deregulation in virtually all cell types in SMA SCOs. They included *CHCHD2*, a mitochondrial protein involved in oxidative phosphorylation and whose upregulation was previously shown in other neurodegenerative conditions to be serving a compensatory protective response against oxidative stress^29,30^. Interestingly, *CHCHD2* has been shown to be subjected to alternative splicing. Overall, SMA SCOs exhibit a disease-associated molecular signature across various MN maturation states, leading to global alterations in multiple molecular programs. Gene ontology analysis defined enriched categories related to metabolic processes, stress response and spinal cord disorders (**Fig. 3G**). Indeed, a significant reduction in neural projections (identified with staining for the heavy neurofilament NF-H/SMI32, a marker gene enriched in MNs) was detected in SMA neurons derived from dissociated SCOs (CTRL 59.17 ± 12.30 µm *vs*. SMA 34.62 ± 8.34 µm, *p* < 0.0001; **Fig. 3H-J**).

### Pathological patterns of activity in SMA spinal cord organoids

Modifications in the electrophysiological activity of spinal MNs have previously been reported in 2D cultures^31^ and pre-symptomatic mouse models of SMA^32^. As in other motor degenerative disorders^33^, this abnormal activity might contribute to the SMA human disease phenotype. However, it is not clear how early in disease progression this dysfunction occurs and how it varies across CNS compartments. For the first time, we exploited high-density multi-electrode array (HD-MEA) technology to investigate the functionality of SCOs derived from SMA patients (**Fig. 4A-C**). Basal recordings of 2-month-old CTRL and SMA SCOs showed spontaneous spiking activity, with significantly lower frequencies in SMA SCOs (CTRL 1.956±0.5354 Hz *vs.* SMA 1.355±0.6807 Hz, p=0.0127; **Fig. 4B, C**). Moreover, SMA SCOs significantly increased their firing frequency more than CTRL SCOs after a glutamatergic stimulus, suggesting an early dysfunction with features of hyperexcitability (CTRL 44.06%±20.03 *vs.* SMA 91.03%±28.42, p<0.0001; **Fig. 4B, C**).

**Figure 4.**
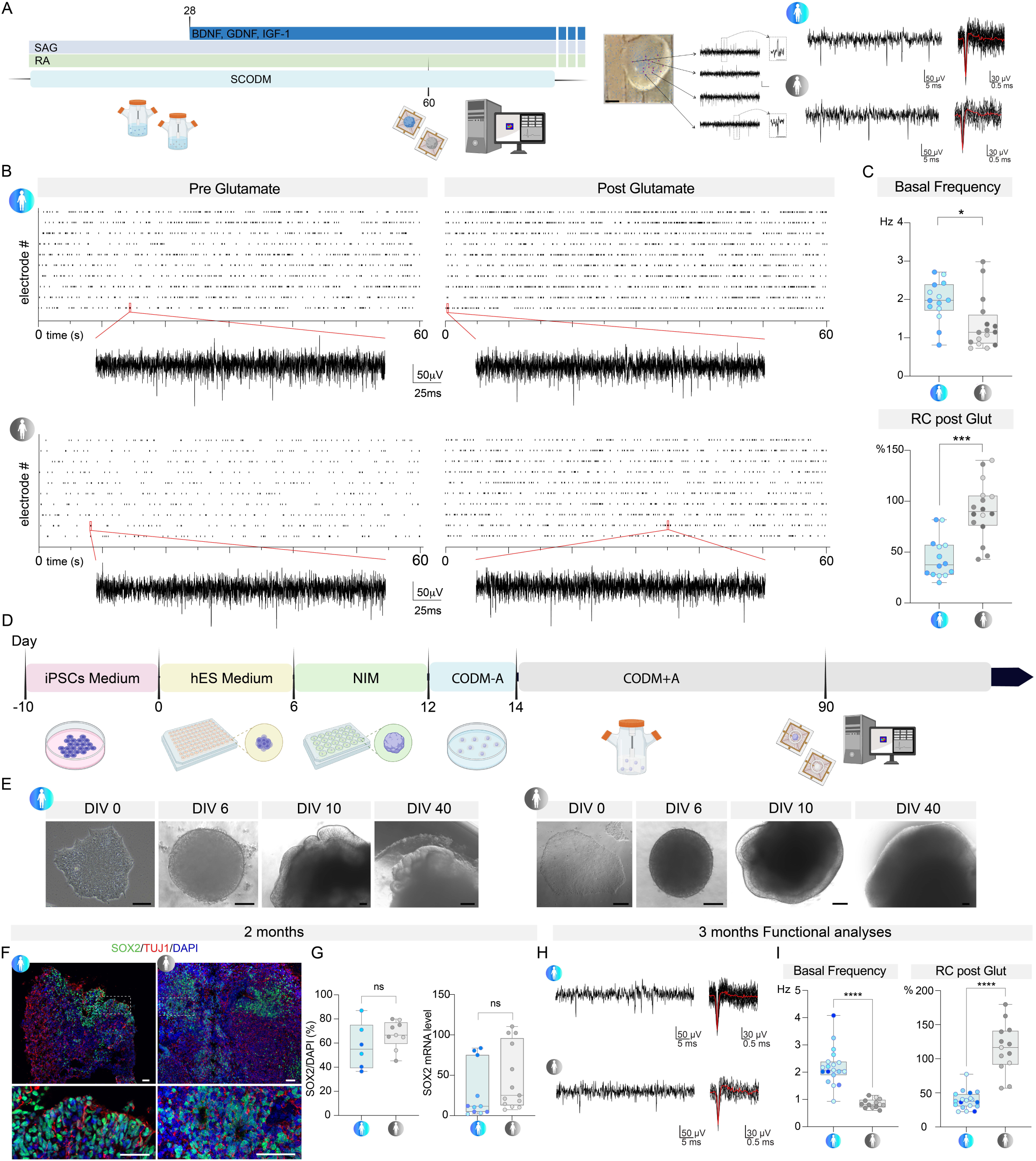
SMA SCOs and cerebral organoids (CrOs) show alterations to electrophysiological activity with features of hyperexcitability. **A**, Schematic of MEA analysis time-line and representative image of SCO on HD-MEA chip with recorded traces showing basal activity. Scale bar: 500 µm. **B**, Raster plot of CTRL and SMA SCOs pre- and post-glutamate perfusion. **C**, Quantification of basal firing activity in CTRL vs. SMA SCOs (unpaired t-test, *p*= 0.0124) and relative change in firing frequency increase after glutamate (glut) perfusion in CTRL *vs*. SMA SCOs (unpaired t-test, *p*< 0.0001). **D**, Schematic of human CrOs generation. **E**, Representative bright-field images of the progression of the differentiation pathway for CTRL and SMA CrOs. Scale bar = 100 µm. **F**, Representative images of SOX2 and TUJ1 staining at 2 mo in CTRL and SMA CrOs. Scale bar: 200 µm. Nuclei were stained with DAPI (blue). **G**, Quantification of SOX2 as a percentage of positive nuclei and analysis of mRNA level in CrOs at 2months. In both analysis the expression of SOX2 does not differ in CTRL and SMA CrOs (Left graph: unpaired t-test, *p*= 0.2449. Right graph: unpaired t-test, *p*= 0.2524). **H**, Representative traces of basal recording of CTRL and SMA cerebral organoids (3 mo). **I**, On the left, basal firing frequency was lower in SMA CrOs than in CTRL (unpaired t-test, *p*<0.0001). On the right, the relative increase in firing frequency after glutamate (glut) perfusion was higher in SMA than in CTRL CrOs (unpaired t-test, *p*<0.0001).

Notably, differentially expressed genes (DEG) analyses revealed, across different clusters, the presence of a set of deregulated genes related to synaptic channels among different neuronal populations, suggesting an overall impact on synaptic function (**Suppl. Fig. 5A**).

To rule out changes in GABAergic transmission, we tested spontaneous firing frequency upon application, at the basal level, of gabazine and strychnine^34^ to block GABA-A and glycine receptors. As expected, an increased spontaneous firing frequency was observed in CTRL organoids (CTRL 44.21% ± 5.626) and no differences were detected in SMA SCOs (SMA 43.75% ± 16.4, *p* = 0.9521; **Suppl. Fig. 5B-C**). This response suggested the presence of active mechanisms of neuronal inhibition in the organoid circuits (CTRL 43.71% ± 36.38 vs. SMA 35.72% ± 19.11, *p* = 0.5455). GAD65 and GABA staining confirmed the presence of inhibitory interneurons in both CTRL and SMA SCOs (**Suppl. Fig. 5B**), as also highlighted by single-cell molecular data. These results support the hypothesis that the observed hyperexcitability response of SMA organoids to glutamate does not imply a reduced inhibitory transmission, and it is more likely to be ascribed to alterations in glutamatergic transmission maturation. Overall, we showed that SCOs display basal firing activity with the potential to respond to an exogenous stimulus and they generate functional circuits. Furthermore, SMA SCOs possess distinct electrophysiological properties, including increased glutamate sensitivity, that can potentially lead to hyperexcitability, a common trait of many neurodegenerative disorders.

### Pathological patterns of activity in SMA brain organoids

SMA has traditionally been considered a lower MN disease; nonetheless, cerebral involvement has been reported in the most severe SMA cases^10^. To begin exploring this component, we employed a protocol for generating 3D brain organoids^14^ from SMA and healthy iPSC lines (**Fig. 4D, E** and **Suppl. Fig. 6**), where a putative secondary effect of spinal cord degeneration was excluded. Both CTRL and SMA cerebral organoids displayed complex morphology, with heterogeneous regions containing neural progenitors (SOX2+), mainly localized within neural rosettes, and neurons (TUJ1+; **Fig. 4F**). We quantified the number of SOX2-positive progenitors across CTRL and SMA brain organoids at both the RNA and protein levels and did not detect a significant difference at the 2-month time point (CTRL 57.45 ± 19.10 *vs.* SMA 67.02 ± 11.54, *p* = 0.2449. **Fig. 4G**). Staining for glial populations (GFAP+) and cortical neurons (CTIP2+ and SATB2+) at 2-months revealed the presence of a variety of cell types (**Suppl. Fig. 6A, B**). As for SCOs, brain differentiation can take place from SMA reprogrammed pluripotent cells and cerebral organoids are able to recapitulate key early steps of brain development.

Interestingly, cerebral organoids displayed abnormal electrophysiological activity like that seen in the SMA SCO. HD-MEA analysis (**Fig. 4H, I**) revealed that SMA cerebral organoids had a significantly lower basal spike frequency compared to CTRLs (2.206 ± 0.6826 Hz SMA *vs.* 0.8497 ± 0.1629 Hz CTRL, *p* <0.0001: **Fig. 4I**). Consistent with the SCO response observed after glutamatergic stimulation, SMA cerebral organoids increased their firing frequency by a significantly higher factor than CTRL organoids (115.3% ± 36.20 SMA *vs.* 39.11% ± 13.21 CTRL, *p* <0.0001: **Fig. 4I**). Moreover, the presence of an active inhibitory neuronal circuit exploiting GABA-A (and likely glycine) receptors was detected (**Suppl. Fig. 6D, E**). In parallel, calcium imaging analyses on whole organoids (**Suppl. Fig. 6F-H**) confirmed that SMA cerebral organoids presented a significantly stronger response to a glutamatergic stimulus compared to CTRL (149,162 ± 135,032 *vs.* 1582 ± 1997, *p* = 0.0232; **Suppl. Fig. 6G, H**).

Our results thus show pathological features in SMA cerebral organoids, demonstrating direct involvement of the disease in the CNS, beyond the spinal cord, and revealing the need to consider additional targets in the clinical therapeutic setting to evaluate the efficacy and the long-term response to treatments.

### Optimized SMN-targeted approach effectively restores cellular and functional alterations

Among the approved therapies for SMA^12^, Nusinersen is based on an antisense oligonucleotide (ASO) acting on *SMN2*, increasing the overall levels of SMN protein. To determine if SMA SCOs are a suitable model for testing the effectiveness of potential therapies, and whether increasing SMN protein levels can correct specific molecular disease signatures, we tested a recently validated ASO for *SMN2* with optimized morpholino chemistry (MO-10-34^35,36^) conjugated with an arginine-rich cell-penetrating peptide (r6-MO). We recently demonstrated that the novel r6-MO can increase drug biodistribution and efficacy in animal models^37^.

Notably, the differentiation phenotype was completely restored by r6-MO treatment (percentage of cells positive for TUJ1: SMA 20.99% ± 8.044, *vs* r6-MO 33.93% ± 7.648, *p* = 0.0003; **Fig. 5B, C**). The administration of r6-MO at day 45 was also able to significantly prevent cell death (percentage of apoptotic nuclei in each organoid at 2 months: untreated 18.16% ± 8.443 *vs.* r6-MO 10.58% ± 3.37, *p* = 0.005; **Fig. 5D, E**), supporting the idea that early intervention could halt the aberrant cellular processes, positively changing the course of the disease.

**Figure 5.**
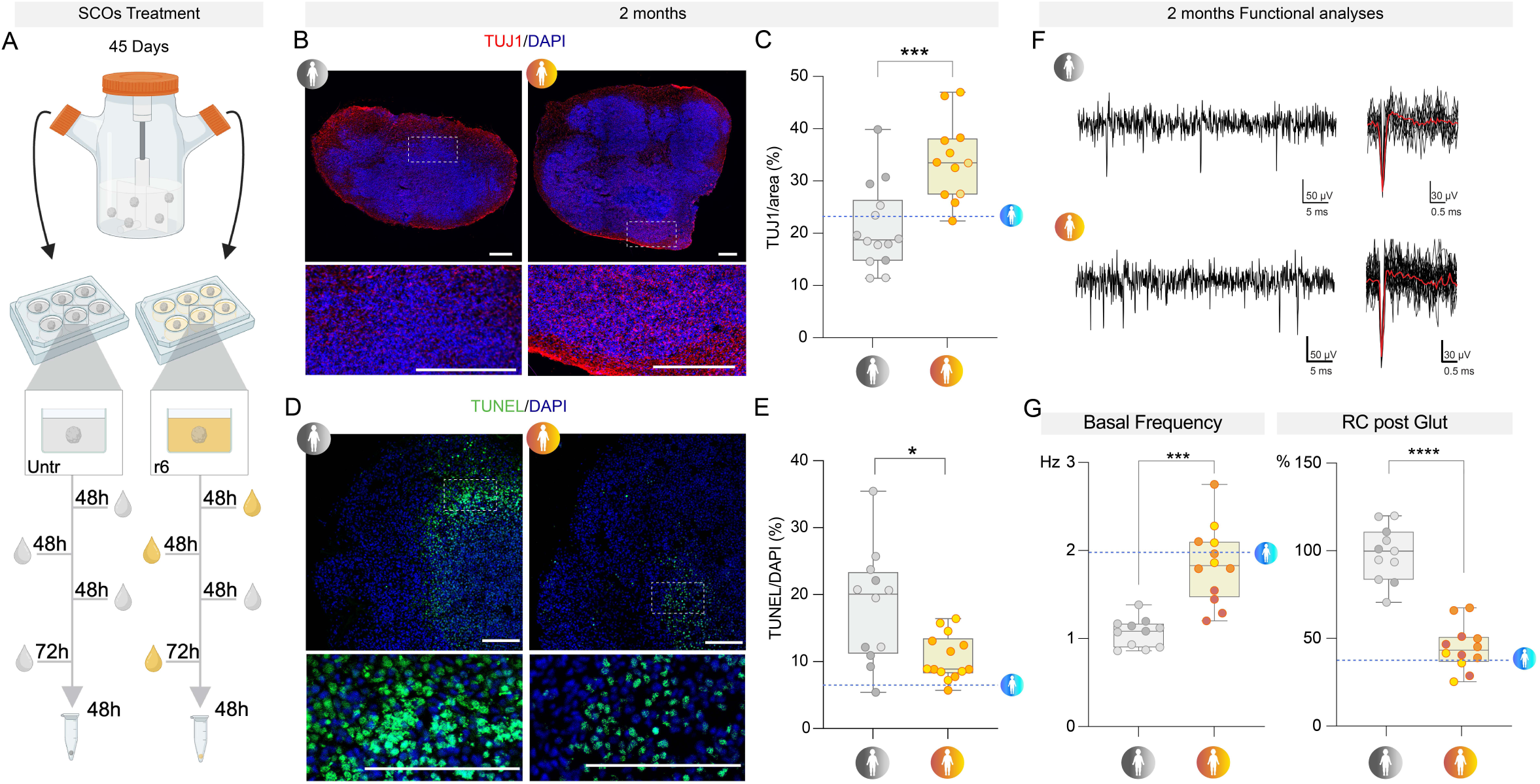
SMA organoids display specific features that are rescued by r6-morpholino. **A**, Schematic representation of r6-morpholino (r6-MO) treatment. SMA SCOs were treated three times: T1 (initial treatment), T2 (48 hours after T1), and T3 (72 hours after the compound was removed and the media was changed). **B**, Representative images of TUJ1 staining of Untr and r6-MO-treated SMA SCOs at 2 mo. Scale bar: 300 µm. Nuclei were stained with DAPI (blue). **C**, TUJ1 expression (intensity/area) is increased in r6-MO–treated SCOs in comparison with untreated SCOs (unpaired t-test, *p*=0.0003). **D**, Representative images of TUNEL staining of Untr and r6-MO-treated SMA SCOs at 2 mo. Scale bar: 300 µm. Nuclei were stained with DAPI (blue). **E**, TUNEL-positive cells were less represented in r6-MO–treated SCOs than in untreated SCOs (unpaired t-test, *p* =0.0050). **F**, Representative traces of basal recording of CTRL and SMA SCOs. **G**, Left panel: quantification of basal firing activity in Untr vs. r6-MO (unpaired t-test, *p* <0.0001). Right panel: relative change in the firing frequency increase after glutamate perfusion (glut; unpaired t-test, *p* <0.0001).

Strikingly, at the functional level, treatment with r6-MO resulted in enhanced basal activity compared to untreated SMA SCOs (r6-MO 1.843 ± 0.4396 Hz *vs*. untreated 1.067 ± 0.1623 Hz, *p*<0.0001; **Fig. 5F, G**), which was paralleled by a reduced response to the glutamatergic stimulus (percentage increase in frequency r6-MO 44.76% ± 12.81% *vs*. untreated 98.31% ± 15.56%, *p*<0.0001; **Fig. 5F, G)**.

These findings reveal that fundamental cellular and functional phenotypes could be rescued by an early specific SMN-increasing therapy across multiple genetic backgrounds. Taken together, these data provide novel insights into the value of optimized ASOs and their impacts on the early treatment of SMA pathology.

### SMA splicing defects are rescued by SMN-increasing therapy

Disease-modifying therapies are currently changing the natural history of SMA, but the specific molecular pathways affected by SMN-targeted approaches and what is their impact on the different cell types along the developmental trajectory are not fully elucidated. We leveraged our novel platform to dissect the molecular drivers responsible for the rescue in the cellular and functional phenotypes. The SMN protein has a well-recognized crucial role in the splicing machinery^38^. Thus, by RNA deep-sequencing, we investigated changes in splicing events in SMA SCOs, and assess the splicing modifications induced by the MO treatment. We analyzed CTRL (n=3 lines, 3 individual replicates), SMA (n=2 lines, 3 individual replicates) and treated SCOs (n=2 lines, 4 individual replicates) after 2 months of differentiation with r6-MO and r6-scrambled-MO as negative control (**Suppl. Fig. 7**).

In SMA vs. CTRL SCOs, an overall transcriptional alteration was confirmed with 666 genes were upregulated and 463 were downregulated (**Suppl. Fig. 7A**). GO analyses revealed enrichment for progressive encephalopathy disorder, while showing a downregulation for synaptogenesis and glutamate receptor signaling (**Suppl. Fig. 7B**). These findings correlate with the alterations in functional activity displayed by SMA SCOs. Since bulk-tissue RNA-seq reveals averaged gene expression patterns across entire organoids, we utilized our single-cell expression data to infer gene modules and understand co-expression in specific cell types. This analysis identified 5 distinct modules with genes enriched in specific cell types: cycling cells (M1: *CDK1*, *CDKN3*, *NUSAP1*, *TOP2A*, *UBE2S*), progenitors (M2: *BTG1*, *EFNB1*, *HES2/4*, *TUBA1C*), astroglia and progenitors (M3: *GABBR2*, *NTRK2*, *PHGDH*, *PTPRZ1*, *SLC1A4*), intermediate progenitors/early MNs (M4: *FOXP4*, *RARA*, *SP1*, *TEAD2/3*, *TCF3*), and mature MNs (M5:*GRIA1/2*/4, *MAPT*, *NEFL*, *SCN2A*) (**Suppl. Fig. 7C**). These data further suggest a pervasive phenotype, involving distinct developmental dynamics, where MNs, along with differentiating cells, are significantly affected by the lack of *SMN1*.

To uncover the molecular mechanisms downstream of r6-MO treatment and elucidate the pathways leading to functional rescue, we compared the transcriptional and splicing landscapes of r6-MO-treated SCOs with those treated with r6-scramble-MO. The r6-scramble-MO treatment served as a negative control, providing a baseline for comparison with r6-MO treatment. Relatively few transcriptional changes were detected after r6-MO treatment compared to r6-scrambled-MO, suggesting that, at least within the considered time window, a global transcriptional change is not required for phenotypic rescue.

In contrast, analysis of pre-mRNA splicing patterns in SMA vs. CTRL, and r6-MO vs. r6-MO-scrambled treated SMA organoids showed substantial differences across conditions (**Fig. 6A, B** and **Suppl. Fig. 7D**). The loss of *SMN1* in SMA patients SCOs (comparing CTRL with SMA organoids) and SMN rescue by r6-MO treatment (comparing r6-scramble-MO with r6-MO) led to multiple significant alternative splicing changes. The most altered events resulting were found as skipped exon (SE), mutually exclusive exons (MXE), followed by intron retention (IR), 3’ (A3SS) and 5’ (A5SS) splice site events, detected with a lower frequency. To assess how r6-MO treatment affects the splicing landscape of SMA cells, we compared pre-mRNA splicing events in r6-MO treated SCOs to those in r6-scramble-MO (baseline control) treated cells. A total of 2614 splicing events resulted to be specifically modified by r6-MO (**Fig 6C**). The most abundant ones were represented by SE, MXE, and IR in similar proportions to the ones observed for the comparison of CTRL *vs* SMA. We could identify 1982 genes specifically associated with the 2614 rescued splicing events. GO enrichment analysis of these genes showed significant enrichment for regulation of chromatin organization, DNA metabolic process, and cell cycle, as well as axon guidance and neurodegeneration (**Fig. 6C**). Importantly, changes in splicing events could be aligned with specific gene modules from the single cell RNA-seq analysis (**Fig. 6D**). We leveraged all the detected rescued spliced genes to compute module score analysis, revealing unique combinations of spliced genes specifically associated with cycling cells (<u>M1</u>: *BRIP1*, *CENPK*, *HMMR*, *TOP2A*, *UBE2C*), progenitors (<u>M2</u>: *ALG3*, *EIF5*, *LTN1*, *MRPL21*, *MRPL13*), progenitors and neurons (<u>M3</u>: *AHI1*, *CEP290*, *KIAA0586*, *VPS13C*, *XPO1*) astroglia (<u>M4</u>: *BMPR1B*, *DDX3X*, *PDGFRB*, *SIRT2*, *SLIT2*) and MN cell types (<u>M5</u>: *GRIA2*, *MAP2*, *SCN1A*, *SCN3B*, *STMN4*) (**Fig. 6D**).

**Figure 6.**
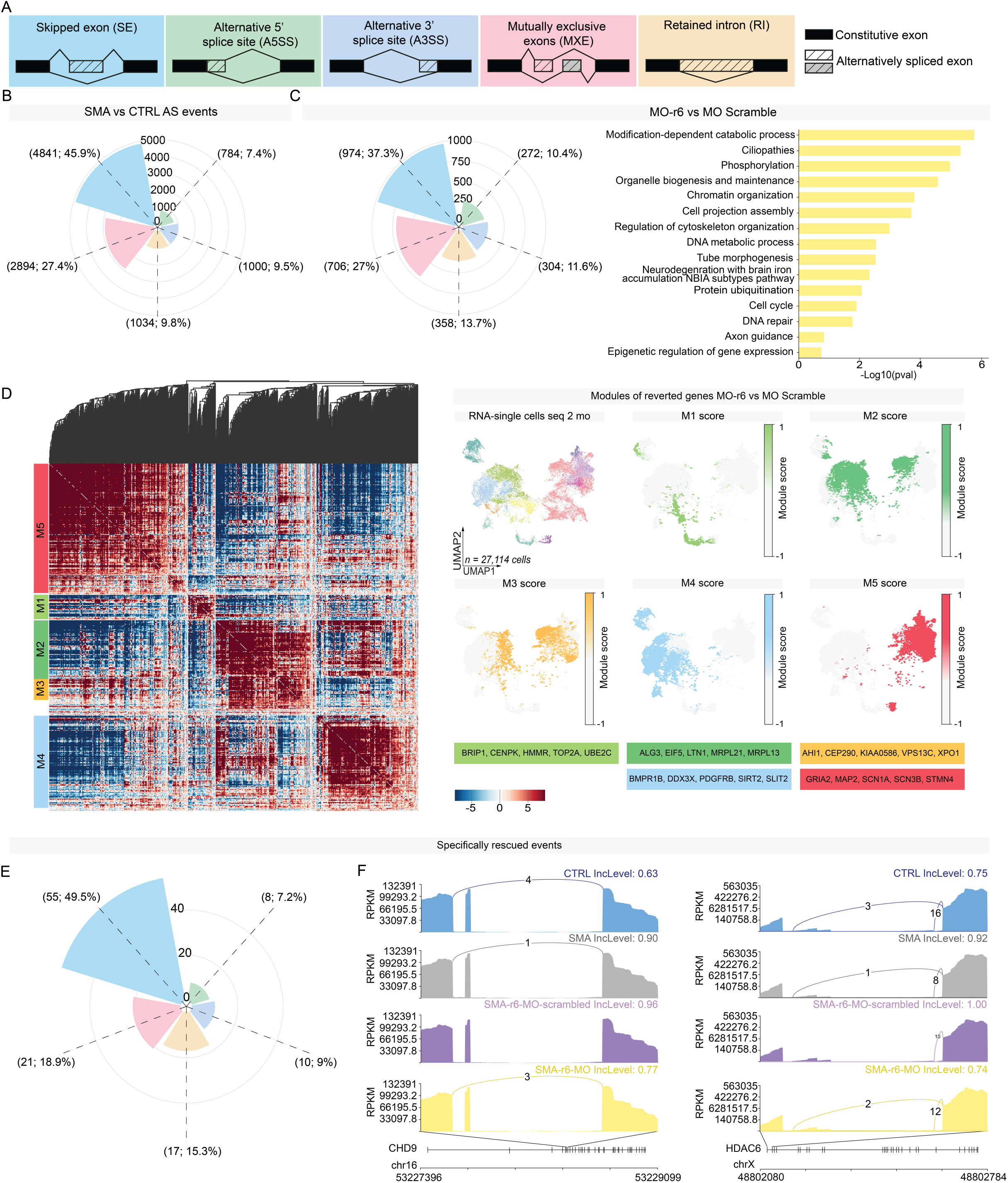
r6-MO treatment modulates splicing event in SMA SCOs. **A**, Schematic of the main classes of alternative splicing events (AS). **B**, Rose plot showing the frequencies of AS events, categorized by major event classes, based on a comparison between SMA *vs.* CTRL. **C**, Rose plot showing the frequencies of AS events, categorized by major event classes, based on a comparison between r6-MO *vs.* MO scrambled treatment in SCOs. Gene Ontology, performed with Metascape, for the 1,982 genes with an associated r6-MO specific splicing event. **D**, Clustermap showing the single-cell co-expression profiles of 907 genes with splicing events specific to r6-MO at 2 months in SMA single-cell RNA-seq, organized into distinct modules. Genes are grouped into modules based on shared co-expression profiles in low-dimensional space. Module scores are highlighted on the UMAPs of individual modules, with key genes highlighted for each module. **E**, Rose plot showing the frequencies of AS events, categorized by major event classes, based on the overlap between the comparison SMA *vs.* CTRL and r6-MO *vs.* MO scrambled. These AS events are represented by those specifically reverted by r6-MO treatment in SCOs. **F**, Sashimi plots display the read distribution and splice junctions for representative exon skipping events in *CHD9* and *HDAC6*, across the four experimental groups.

Our findings indicate lineage-specific splicing alterations downstream of r6-MO treatment. To assess whether these changes restore the splicing landscape of specific genes toward control levels, we compared splicing events related to the SMA phenotype vs. CTRL with those induced by r6-MO (carrying along the scramble-MO as specific control of the treatment). To do that, we intersected splicing events induced in SMA vs. CTRL and r6-MO vs. r6-scramble-MO conditions. This integrated analysis revealed a distinct set of splicing events *rescued* upon r6-MO treatment (**Fig. 6E**). Notably, genes outside this category (which did not fully recover their splicing pattern by the r6-MO treatment) were linked to broader pathophysiological processes, further suggesting a systemic involvement beyond spinal cord diseases. Early r6-MO treatment restored splicing isoforms in 98 genes, which were identified as novel top candidates in our bioinformatics analysis. These genes likely play a crucial role in alleviating the phenotypic defects of SMA, with several chromatin remodeling enzymes, *ARIH2, ATG4A/B, CHD9*, and *HDAC6*, emerging as key players. Over time, these targets may help re-establish the molecular profile of both progenitors and neurons, thus contributing to the observed cellular and functional rescue.

Among the rescued genes, we highlighted *CHD9* (Chromodomain Helicase DNA-binding protein 9), a key regulator of the cell cycle in embryonic stem cells (**Fig. 6F, sashimi plot**), and *HDAC6* (Histone Deacetylase 6), which is critical for MN development and function through its roles in axonal transport, autophagy promotion, and neuroprotection. These findings suggest that rescued splicing events impact diverse cellular and molecular processes crucial for progenitor and neuronal development, driving the observed functional recovery following r6-MO treatment.

Interestingly, although these ‘reverted’ genes lacked class-specific expression in our single-cell dataset, they were significantly enriched in ‘SMA unmatched clusters’, comprising progenitors and neurons without clear correlates in control lines (**Suppl. Fig. 7E, F**). This confirms that an early splicing defect – caused by *SMN1* deficiency –impacts the molecular dynamic of spinal cord development, subverting the physiological trajectory of neuronal maturation and function.

Overall, these results point to a scenario in which the restoration of proper splicing landscape of genes associated with spinal cord development is among the key molecular events occurring upon MO treatment in SMA cells. This also suggests that in SMA organoids aberrant splicing dynamics impair progenitor and neuron development and function, and that r6-MO induced splicing correction thus drives and/or contributes to the rescue of their cellular and functional phenotypes.

## Discussion

Stem cell–based models of the human CNS provide an opportunity to unravel processes of human biology that could not otherwise be studied experimentally^42–44^. Cerebral organoids have been proposed as a platform to study human brain development and identify disease phenotypes^43,45,46^. However, fewer descriptions of spinal cord-like organoids have been reported in the literature^16,47,48^, and this area of research deserves further investigation in the context of MN diseases, and neurological disorders with urgent unmet therapeutic needs. Here, we leveraged the organoid technology to generate SCOs and cerebral organoids from iPSCs of healthy subjects and SMA type 1 patients to further our understanding of SMA and test innovative optimized therapies.

The advent of disease-modifying therapies for SMA has changed the scenario of clinical practice and offered new expectations to patients, caregivers, and healthcare providers. Current evidence demonstrates that early treatment is essential for clinical efficacy. Thus, increased knowledge of the prenatal and pre-symptomatic stages of the disease is crucial to optimize therapeutic strategies. Even though SMA is a genetic disorder often with onset in newborns, very few studies have analyzed the prenatal stages of the disease, when SMN function is critical^8^, and no molecular data on human SMA spinal cord at single-cell resolution have been reported thus far, exploring how different cell populations are affected. Prenatal MN cell death and dysfunction might condition the onset and severity of the disease postnatally and set the therapeutic window. Still largely unexplored is whether SMN deficiency impairs early neuronal development and thereby triggers neuronal death. In this perspective, SMA organoids represent optimal substrates to begin addressing the link between key early pathological features. Here, we demonstrated that both CTRL and SMA SCOs harbor a spectrum of spinal neuronal domains recapitulating the *in vivo* complexity of neuronal populations within the human developing spinal cord (**Fig. 1**). Moreover, we showed that early differentiation programs are affected in SMA spinal organoids (**Fig. 2**) even before cell death becomes prominent. We identified the presence of a higher number of cells within the early developmental states in the SMA SCOs (**Fig. 2**). These data, together with the pervasive molecular alterations observed in both progenitors as well as neuronal types at the same stage when cell death increases (2mo, **Fig. 3**), point towards an important developmental component in SMA pathogenesis and suggest that an aberrant developmental trajectory might negatively impact on the differentiation and survival of distinct neuronal subtypes. The developmental component of a neurodegenerative disorder such as Huntington’s disease has been recently studied^49–51^ suggesting that this hypothesis can be investigated also in SMA.

The observation of reduced neurite length in SMA SCOs relative to CTRL demonstrates that the 3D model can reproduce key disease features of SMA (**Fig. 3**). Intriguingly, early neuronal differentiation defects and apoptosis were reproduced in SMA SCOs, suggesting that concomitant defects in differentiation and increased degeneration could contribute to SMA pathogenesis.

Interestingly, when we looked at differentially expressed genes, we observed that a large fraction of them was detected in other cellular subgroups other than MNs. This data suggests that, despite defects of spinal MN function being the central feature of SMA pathology, the disease also has a broader impact. A growing body of evidence suggests that SMA is a multifactorial disease in which the interaction of different cell types and disease mechanisms progressively leads to MN dysfunction and death^11,52^. Magnetic resonance imaging has revealed that severe SMA patients who survive beyond 1 year of age exhibit progressive and diffuse brain lesions^10^. Prenatal spinal cord alterations have been reported in animal studies^52^, and recently in humans^9^. A very recent study demonstrated functional alterations of the motor cortex and cerebellum in symptomatic SMA mice^53^. Herein, we demonstrated that SMA cerebral organoids show impaired neuronal activity (**Fig. 4**). This wider impact requires further characterization and needs to be considered for the effective clinical management of SMA patients. Indeed, the generation of spontaneous and evoked activity in both SCOs and brain organoids is impaired (**Fig. 4**), demonstrating the biological relevance of the model and the direct impact that developmental alterations and/or plasticity events might cause on neuronal wiring. Neuronal hyperexcitability has been described in the context of SMA^31,54^ and ALS^55^, associated with degeneration of both upper and lower MNs^56,57^. However, it is not clear how early in SMA progression this dysfunction arises and how it varies across CNS compartments. SMA SCOs showed decreased basal activity and hyperexcitability early on, likely contributing to the disease phenotypes. Furthermore, we reported an SMN-dependent deregulation of glutamate channel genes in multiple neuronal populations, which may point to intrinsic transcriptional changes in SMA neurons related to the observed hyperexcitability.

The last decade has witnessed the development of a growing number of novel therapies for SMA. However, the FDA/EMA-approved treatment nusinersen^58^, an ASO able to increase SMN expression by splicing correction of *SMN2*, requires administration by periodic lumbar puncture. Permeability of the blood–brain barrier and tissue distribution of ASO delivery can be fostered by CPPs, small cationic molecules with carrier capacity for macromolecular compounds. This strategy offers improved systemic distribution, including upon intravenous injection, in several disease models^36^ and was proposed in the context of Duchenne muscular dystrophy^59^. In this current study, for the first time, we tested the same conjugate in human tissue in SMA SCOs and rescued SMA pathological hallmarks (**Fig. 5**).

We exploited deep sequencing bulk RNA-seq to globally analyze the altered splicing events in SMA organoids (**Fig. 6**). This allowed us to evaluate how splicing changes were present in different genetic backgrounds and how the SMN correction had a pervasive effect, independently of the patient-specific background, at the level of different cell types. Indeed, using single-cell RNA-seq data as a reference, we were able to assess how these rescued targets were expressed by different cell populations in varying developmental states and to uncover important molecular events related to spinal cord development in SMA pathogenesis. For the first time, here, we identified splicing alterations associated with SMA phenotype, whose correction might contribute to the rescue of key-pathological dysfunction.

This not only opens new avenues of investigation to fully dissect the therapeutic mechanisms of action on the multifaceted aspects of SMA, but also confirms these novel models as suitable tools to improve the current therapeutic strategies in the clinic, and even expand to more personalized medicine in the future. Altogether, our study suggests that SMN is a key molecule during prenatal spinal cord and brain development, meaning that SMA therapeutics will be most effective if delivered during the pre-symptomatic phase of the disorder, namely peri- or even prenatally. Further studies to analyze the multisystemic impact of SMN deficiency during development will be pivotal in defining the optimal therapeutic window and determining the efficacy of therapies on early developmental defects.

## Materials and methods

### iPSC reprogramming and culture

The studies involving human samples were conducted in accordance with the ethical standards of the Declaration of Helsinki, national legislation, and institutional guidelines. Human fibroblast cell lines were obtained from Eurobiobank with informed consent, as approved by the ethical committee at Fondazione IRCCS Ca’ Granda Ospedale Maggiore Policlinico and University of Milan (0004520). Fibroblasts derived from skin biopsies were reprogrammed into iPSCs using the CytoTune iPS 2.0 Kit (CytoTune-iPS Sendai 2.0 Reprogramming Kit, Invitrogen) and maintained in E8 (Essential 8™ Medium, Gibco, A1517001) medium at 37°C in 5% CO_2_^60^.

### Generation of cerebral organoids

Cerebral organoids were generated by adapting a previous protocol^14^. Selected iPSCs were harvested using Accutase (Life Technologies, A1110501) and approximately 9,000 iPSCs were plated in each well of an ultra-low attachment 96-well plate (Corning, CLS7007) in pluripotent stem cell media containing low recombinant human FGF-basic (4 ng/ml, Peprotech, 100-18B) and Y-27632 (50 μM ROCK inhibitor Calbiochem, 13624S). On day 6, obtained embryoid bodies (EBs) were transferred to low adhesion 24-well plates (Corning, CLS3473) in neural induction media (NIM) containing DMEM/F12 (Gibco, 11320033), 1% N2 supplement (Gibco, 17502001), 1% Glutamax (Life Technologies, 35050061), 1% MEM-NEAA (Gibco, 11140035), and 1 µg/ml heparin (Sigma, H3149). On day 12, the EBs were transferred to droplets of Cultrex® Basement Membrane Matrix Type 2 (BME type 2, Trevigen, 3532-001-02) and grown in differentiation media containing a 1:1 mixture of DMEM/F12 and Neurobasal (Gibco, 21103049) with 0.5% N2 supplement, 1% B27 supplement without vitamin A (Gibco, 12587010), 2-mercaptoethanol (ThermoFisher 21985023), 0.025% insulin (Sigma, I9278), 1% Glutamax, and 0.5% MEM-NEAA. After 2 days, the droplets were transferred to a spinning bioreactor containing the same media except for B27 without vitamin A, which was replaced with B27 with vitamin A (Gibco, 17504044). The speed of the spinning bioreactor was gradually increased up to 45 rpm.

### Generation of spinal cord organoids (SCOs)

To generate SCOs, the same procedure described for human cerebral organoids was applied from day 0 to day 2. On day 2, half of the medium was changed to fresh medium containing a final concentration of 4 ng/ml bFGF, 50 μM Y-27632, 3 μM CHIR 99021 (GSK3 inhibitor, Sigma-Aldrich, SML1046), 2 μM SB-431542 (Activin/BMP/TGF-β pathway inhibitor, Sigma-Aldrich, S4317), and 0.2 μM LDN-193189 (inhibitor of the bone morphogenetic protein [BMP] pathway, Stemcell Technologies, 72149). On day 4, the medium was replaced, adding the same small molecules (excluding bFGF and Y-27632). On day 6, the EBs were transferred into ultra-low attachment 24-well plates and cultured in NIM plus 100 nM retinoic acid (RA, Sigma-Aldrich, R3255). On day 12 EBs embedded in Cultrex and cultured with SCO differentiation media (SCODM, composed of 1:1 DMEM/F12 and Neurobasal (Gibco, 21103049) with 0.5% N2 supplement, 1% B27, 2-mercaptoethanol (ThermoFisher 21985023), 0.025% insulin (Sigma, I9278), 1% Glutamax, and 0.5% MEM-NEAA). 1 μM Smoothened Agonist (SAG, Calbiochem, 364590-63-6) was added to the media. On day 14, SCOs were transferred to spinning bioreactors and cultured with SCODM supplemented with RA and SAG. On day 28, concentrations of RA and SAG were reduced to 50% and neurotrophic factors BDNF (Peprotech, 450-02), GDNF (Peprotech, 450-10), and IGF-1 (Sigma Aldrich, 100-12) were added at 10 ng/ml each.

### Immunohistochemistry analysis

Organoids were fixed with 4% paraformaldehyde (PFA) for 15 min at room temperature (RT) and subsequently placed in 30% sucrose/PBS overnight at 4°C. The next day, they were embedded in 10%-7.5% gelatin/sucrose, frozen in dry ice, placed at -80°C overnight, and cryosectioned into 20-μm sections. Organoid sections were permeabilized for 1 h in PBS with 0.3% Triton-X (Sigma) and 10% donkey or goat serum (Jackson ImmunoResearch) at RT.

2D cultures were fixed with 4% PFA at RT for 5 min and permeabilized in PBS with 0.2% Triton-X (Sigma) and 10% donkey or goat serum (Jackson ImmunoResearch) for 1 hour at RT. Primary antibody diluted in fresh permeabilization solution was added overnight at 4°C. The following day, secondary antibodies (AlexaFluor 488, 555, 568, 594, and 647 conjugates, 1:1000; Invitrogen) were added with DAPI (1:500; Sigma-Aldrich, D9542) in PBS for 90 min at RT. DAKO mounting media (Agilent, S3023) was used to mount the coverslips. TUNEL assay was performed using a DeadEnd™ Fluorometric TUNEL System (Promega, G3250). Confocal images were obtained using a Leica SP8 white laser inverted confocal microscope (Leica) and analyzed using Fiji image-processing software.

### Image quantification

To evaluate the number of SOX2^+^ cells and the percentage of TUJ1-positive area, 10× mosaic images of whole SCOs sections were acquired using an Inverted Microscope DMi8 and the Leica LASX software platform. 20× mosaic images of whole brain organoid sections were imaged using Zeiss Axio Microscope and the Zen Light software platform. To compute the number of SOX2^+^ cells, a custom-written Cell Profiler^61^ pipeline was designed, and a custom-designed ImageJ/Fiji^62^ software image macro was applied to evaluate the proportion of TUJ1^+^ area (as %). For TUNEL quantification, images were acquired by an inverted Nikon-Crest multimodal spinning-disk confocal microscope. Quantification of nuclei (DAPI) and TUNEL-positive cells was performed with an ad-hoc designed analysis pipeline for thresholding and segmentation using the GA2 module in Nis-Elements v5.21 software (Nikon-Lim Instruments). Regarding neurofilament tracing, images previously acquired with a Leica SP8 white laser inverted confocal microscope were analyzed using Fiji image-processing software. The NeuronJ plug-in was used for the tracing and analysis of neurite length; only filaments that both originated and terminated within the field of view were traced. Filaments extending beyond the field of view or those for which the origin could not be determined were excluded from consideration.

### Quantitative real-time PCR (qRT-PCR)

Total RNA was extracted using the ReliaPrep RNA Cell Miniprep System (Promega, Z6010). The RNA concentration was tested using Nanodrop (Thermo Fisher Scientific). Reverse transcription was performed using SuperScript IV VILO Master Mix (Thermo Fisher Scientific). 150 ng of template total RNA were retro-transcripted. This protocol has been used to process bulk RNA samples. The expression level of candidate genes was assessed by: i) TaqMan quantitative analysis on the 7500 Real-Time PCR System and 7900HT Fast Real-Time PCR System (Applied Biosystem), GAPDH was used as a reference gene; ii) Powertrack SYBR (Thermo Fisher Scientific) quantitative analysis on the ViiA 7 Real-Time PCR System (Thermo Fisher Scientific), B-ACTIN was used as a reference gene.

### Calcium imaging analysis

Fluo-4 AM (Thermo Fisher, F14201), a cell-permeant assay, was used to image the spatial dynamics of Ca2+ signaling in cerebral organoids. Organoids were placed in 8-well chambers (Starsted), one in each well, and washed with PBS without Ca2+ and Mg (Euroclone, ECB4004). A final concentration of 10 μM Fluo-4 was diluted in Opti-MEM (Gibco), added to the wells, and incubated at 37°C for 30 min. Organoids were then washed three times with PBS without Ca2+ and Mg and incubated with pure Opti-MEM for 15 min at 37°C. Calcium flux was visualized and acquired using an LED-illuminated automated spinning-disk video-microscope (Nikon Ti equipped with CREST-Optics X-LightV2 head and an Andor DU888 EM-CCD camera for ultra-fast acquisitions). After 10 min recording, glutamate solution (100 μM in Opti-MEM, Sigma-Aldrich) was added and changes were recorded for an additional 10 min. Time-lapse videos were recorded every 5 s for a total of 20 min.

### Electrophysiology

High-density multielectrode array (HD-MEA) was used to record the extracellular spontaneous activity of both SCOs and cerebral organoids. The device consists of 4096 planar electrodes measuring 21 μm × 21 μm, covering an area of 2.67 mm × 2.67 mm (Biocam X with Biochip Arena, 3Brain AG, SwissSwitzerland). Organoids were gently positioned on the chip and fixed with a platinum anchor and with a nylon mesh. Before and during recordings, organoids were submerged in Kreb’s solution with the following composition (mM): 120 NaCl, 2 KCl, 1.19 MgSO4, 26 NaHCO3, 1.18 KH2PO4, 11 glucose, 2 CaCl2, oxygenated with a mix of 95% O2/5% CO2 to obtain pH 7.4. Organoids were recorded under control conditions for 15 min, after addition of 100 μM of glutamate (L-glutamic acid, Sigma-Aldrich) for an additional 15 min, and after adding the combination of 100 μM glutamate, 10 μM gabazine (SR-95531, Abcam), and 1μM strychnine hydrochloride (Sigma-Aldrich) for an additional 15 min.

Brainwave X (3Brain AG) software was used to record the electrophysiological activity at the sampling frequency of 17 kHz. Brainwave 4 software (3Brain AG) was used to analyze the traces offline. The precise time spike detection (PTSD) algorithm was used to detect the spikes (parameters used: positive and negative peak, peak duration <1 ms, refractory period >1 ms, and detection threshold >7 times the standard deviation of the noise). Spike sorting was performed using a principal component analysis with K-means (silhouette method) considering a maximum three clusters per electrode and three spikes per cluster. The Pakhira-Bandyopadhyay-Maulik (PBM) index was used for validating clustering results^63^. A final checkout was performed by the researcher by re-analyzing the signals and removing any confounders. Only channels showing stable activity during the control period (without spike frequency fluctuations exceeding 20% spike frequency) were taken into consideration. The number of channels per organoid considered for the analysis was comparable between control (SCO 13.6 ± 0.6; cerebral 13.8 ± 2.1), SMA (SCO 15.2 ± 0.7; cerebral 15.4 ± 0.9), and MO-treated SMA (SCO 12.8 ± 1.3) organoids. The percentage change in basal firing frequency was calculated by comparing the average firing frequency in the last 10 minutes before glutamate addition with the interval between 5-15 minutes after glutamate addition.

### Morpholino treatment

We used a morpholino (MO)10-34 sequence targeting the SMN2 ISS-N1 region downstream of exon 7 that was previously described by our lab^35^. We used MO alone and conjugated with r6 peptide (r6-MO)^37^. As negative control, we used a custom-made r6-MO with a scrambled sequence. To assess biodistribution, we used a biotinylated r6-MO that was added to the culture media as described below.

All MO oligomers were synthesized by Gene Tools. We treated SMA SCOs with growing concentrations of MO or r6-MO. The solution containing 2.4 nmoles MO or r6-MO was added directly to each well and incubated for 48h. Afterward, the culture medium was changed and fresh aliquots of 2.4 nmoles MO or r6-MO were added for an additional 48 h. The medium was then replaced with fresh medium for 72 h. Lastly, the organoids were treated again with 4.8 nmoles of MO or r6-MO for 48 h.

### Western blot analysis

Western blot analysis was performed as described previously^35^. Briefly, samples were sonicated in lysis buffer supplemented with protease and phosphatase inhibitors (Pierce Rockford, IL, USA) on ice for 10 min. Approximately 5 μg of protein was separated by 12% SDS-PAGE and electrophoretically transferred to a nitrocellulose membrane. The membranes were incubated with anti-SMN (1:1000, BD) and anti-actin (1:1000, Sigma, St. Louis, MO, USA) antibodies. After labeling with peroxidase-conjugated secondary antibody (Life Technologies), the proteins were detected using a chemiluminescent substrate (Amersham, Pittsburgh, PA, USA). Signal was detected using LI-COR Odyssey 9120. Densitometry analysis was performed using Image Studio Lite ver. 5.2.

### 2D culture of cells from dissociated organoids

SCOs were centrifuged at 300*g* for 5 min and trypsin-EDTA was added before transferring organoids in the water bath at 37°C for 5 min. The organoids were then mechanically dissociated, and a double volume of horse serum (Euroclone, ECS0091L) was used to inactivate the trypsin before centrifugation. The supernatant was discarded, and SCO differentiation media was employed to resuspend the pellet. Approximately 50,000 cells were seeded in 24-well plates with coverslips previously coated with poly-L-ornithine (Sigma-Aldrich, P3655) and laminin (Sigma-Aldrich, L2020). SCO differentiation media supplemented with RA, SAG, and growth factors at the same concentration of differentiation day 28, was used for the maintenance of organoid-derived cells.

### Whole mount staining

Organoids were washed in PBS and then fixed in 4% PFA on a spinning wheel at 4°C overnight. The day after, five PBS washes (10 min each) were performed on the spinning wheel at RT. Organoids were permeabilized with 0.25% Triton X-100 in PBS for 15 min. To prevent non-specific binding, organoids were immersed in PBS with 10% FBS and 0.5% serum. Primary antibodies were changed three times (after 2 h, after 12 h, and after 5 h) and diluted in PBS with 5% FBS, 0.5% serum of the same species as primary antibodies, and 0.1% Triton X-100. Secondary antibodies were diluted in the previously described solution for primary antibodies and incubated overnight at RT on the spinning wheel. Images were acquired using Nikon’s A1 R MP+ multiphoton microscope by the Alembic Facility (IRCCS Ospedale San Raffaele).

### SCO cell dissociation for scRNA sequencing analysis

Organoids were dissociated using an optimized protocol based on the Worthington Papain Dissociation System Kit (Worthington Biochemical). Single cell suspensions were collected on ice-cold PBS (without calcium and magnesium) with 0.04% BSA (Sigma Aldrich) at 1×10^6^/ml as counted by an automatic cell counter (Countess II, Thermo Fisher).

### Single-cell RNA-sequencing

Approximately 8000 cells from each sample prepared as described above were loaded into one channel of the Single Cell Chip B using the Single Cell 3’ v3 reagent kit (10X Genomics) for gel bead emulsion generation in the Chromium system. Following capture and lysis, cDNA was synthesized and amplified for 14 cycles following the manufacturer’s protocol (10X Genomics). The amplified cDNA (50 ng) was then used for each sample to construct Illumina sequencing libraries. Sequencing was performed on the NextSeq550 Illumina sequencing platform following the 10x Genomics instructions for read generation. A sequencing depth of ∼45,000 reads/cell was obtained for each sample.

#### Identification of differentially expressed genes between 2mo CTRL and SMA SCOs Data analysis

Raw sequencing data (bcl-files) were converted to fastq files using the Illumina bcl2fastq tool integrated into the CellRanger (10X Genomics) suite (v=3.0.2). The CellRanger analysis pipeline was used to generate a digital gene expression matrix starting from raw data. Reads were aligned to the human GRCh38 reference genome, annotated, and counted with gene annotations from Ensembl version 93. The CellRanger count module was used to map reads with default settings, and the sequence length was set to r1-length=28 and -r2-length=55. At least 90,000 reads per cell were produced. For further preprocessing and analysis, we used the Scanpy^64^ single-cell analysis toolkit (v=1.9.6, pandas **v=**2.2.0, and numpy v=1.26.3). Doublets were identified and removed with Scrublet^65^ (v=0.2.3). The threshold at doublet scores was automatically set by the Scrublet algorithm and was used individually for each sample. Quality control metrics, including mitochondrial and ribosomal content, were calculated using sc.pp.calculate_qc_metrics function, and then the object was filtered for the outliers cells containing less <10% mitochondrial and <5% ribosomal transcripts. Further, cells were filtered using a sample-specific median absolute deviation (MAD) threshold of “n_genes_by_counts”. After filtering, the data were concatenated using sc.AnnData.concatenate, annotated with sample names and genotypes. We used scVI-tools^66^ (v=1.1.0) to mitigate batch effects where each genotype and sample was treated as a batch. The AnnData object was first transformed by copying counts to adata.layers["counts"] and identifying 2000 highly variable genes using sc.pp.highly_variable_genes with the Seurat V3 flavor, considering the “Sample” batch key. The transformed data were prepared for scVI analysis by setting up AnnData with categorical covariates “Sample” and “genotype”, and continuous covariates “pct_counts_mt”, “pct_counts_ribo”, and “total_counts”. The scVI model was initialised with scvi.model.SCVI and trained using default parameters. The latent representation was extracted using model.get_latent_representation and stored in adata.obsm["X_scvi"]. Dimensionality reduction was performed by constructing a KNN graph with sc.pp.neighbors using the latent representation (X_scvi) and running UMAP with min_dist=0.2. Clustering was executed at resolutions 0.25, 0.3, 0.5, 0.7, and 1.0 using the Leiden algorithm^67^ (sc.tl.leiden), and the results were visualized with UMAP plots (sc.pl.umap) displaying cluster assignments, sample information, and genotype. After integration, the data were reverted to all genes using adata.raw.to_adata(). Raw counts were normalized using sc.pp.normalize_total(adata, target_sum=None, inplace=False) and log-transformed with sc.pp.log1p as suggested in the recent sc-RNA-seq data transformation benchmark study^68^. The final processed data were saved for downstream analysis. Gene set scores were calculated using Scanpy’s sc.tl.score_genes function. To find differentially expressed genes we applied the Wilcoxon rank-sum test on the normalised counts between all found clusters (resolution=0.7), the top 50 genes sorted by the p-value and log-fold change were considered for the biological interpretation.

#### Differential abundance testing

We performed differential abundance testing using the Python version of Milo^24^ (v=0.1.1). Neighborhoods were constructed with a proportion of 0.1 using milo.make_nhoods. Neighborhood counts were computed with milo.count_nhoods using the sample column “Sample”. Both genotypes (CTRL and SMA) were encoded as continuous and categorical variables. Differential abundance testing was conducted using milo.DA_nhoods with a design formula based on genotype. A neighborhood graph was built using milopy.utils.build_nhood_graph and visualized with milopl.plot_nhood_graph, highlighting the spatial false discovery rate (FDR) at 1%. Neighborhoods were annotated with cell types using milopy.utils.annotate_nhoods and the color palette from Leiden clustering was mapped to the neighborhoods.

#### Pseudo-bulk differential gene expression analysis between SCOs genotypes

To test for differentially expressed genes between genotypes for a particular cell identity (at least 25% of cell proportion for each genotype), we leveraged the PyDESeq2^69^ (v=0.4.4) and the Python implementation of decoupleR^70^ packages (v=1.6.0). Pseudo-bulk profiles were generated using the decoupler.get_pseudobulk function with a minimum of 10 cells and 1000 counts per group. The pseudo-bulk data were normalized, log-transformed, scaled, and principal component analysis (PCA) was performed. Associations between metadata and principal components were computed using decoupler.get_metadata_associations, and the results were visualized with decoupler.plot_associations. Data were subset by cell type, and differential expression analysis was performed on selected cell types. Genes were filtered by expression with a minimum count of 10 and a minimum total count of 15. The DESeq2 object was constructed with the design factor ‘genotype’ and reference level ‘CTRL’. Log fold changes (LFCs) were computed, and contrasts between SMA and CTRL genotypes were extracted. Differentially expressed genes were filtered by adjusted p-value (< 0.05) and absolute LFC (> 0.5). The results then were visualised in the form of complex heatmaps and volcano plots with custom R scripts. The lists of differentially expressed genes of selected cell types was analyzed using the Ingenuity Pathway Analysis (IPA - Qiagen Ingenuity Systems) to compute canonical pathways and biological functions enrichment.

#### Trajectory analysis

We performed trajectory analysis on the integrated data using Palantir^71^ (v=1.3.3) and scFates^25^ (v=1.0.6) to model cellular differentiation and progression. scVI (adata.obsm["X_scvi"]) projection was used to run diffusion maps with Palantir (palantir.utils.run_diffusion_maps), generating a multiscale diffusion space with 10 eigenvectors (palantir.utils.determine_multiscale_space). The SimplePPT approach was applied to compose a principal graph of 100 nodes on the multiscale diffusion space embeddings with scFates function scf.tl.tree(adata, method="ppt", Nodes=100, use_rep="X_palantir", device="cpu", seed=42, ppt_lambda=200, ppt_sigma=sig, ppt_nsteps=100). The cycling cells node population was initialized as the root and was chosen based on previous biological knowledge. Then, from the root, the cells were projected onto the principal graph and pseudotime values were calculated with the scFates function “scf.tl.pseudotime with n_jobs=20, n_map=100, seed=42”. Based on the calculated pseudotime and principal graph, the dendrogram was generated with “sc.fates.tl.dendrogram()”. The data were split by genotype into control (CTRL) and SMA groups. Separate dendrograms for CTRL and SMA were plotted using scf.pl.dendrogram. Detailed pseudotime distributions were visualized using KDE plots (seaborn.kdeplot) to show the distribution of pseudotime values across different cell types for both CTRL and SMA groups.

#### Regulons inference and trajectory analysis of motoneurons lineage

To infer the co-expression modules between transcriptional factors and putative target genes in MN cell lineage we used the pySCENIC pipeline^28^. We performed independent pipeline runs for each SCOs genotype data object with the Sungularity container. At the first step of regulons inference, the pipeline relies on a machine learning model that fits the expression patterns of genes to expression patterns of TFs by the GRNBoost2^72^ algorithm. For this part, we used the command line tool (pyscenic:0.12.1 arboreto_with_multiprocessing.py) with the default parameters. Next, the output fits were used to predict the potential regulons and direct-binding targets that are then refined through TFs motifs enrichment analysis with (pyscenic:0.12.1 pyscenic ctx) with “–-maks dropout” parameter. Lastly, cells were scored for each regulon activity in SMA and CTRL objects respectively via the AUCell method^73^ with (pyscenic:0.12.1 pyscenic aucell). Obtained regulons activities scores and metadata were used in the secondary analysis for visualisation and trajectory analysis. We performed trajectory analysis on SCOs focusing on MN linage using Palantir and scFates. The SCOs data object was loaded, and specific clusters unrelated to the MN lineage were removed. Diffusion maps were computed to generate a multiscale diffusion space with four eigenvectors. The principal tree was constructed using scf.tl.tree with parameters method="ppt", Nodes=200, ppt_lambda=100, ppt_sigma=sig, ppt_nsteps=200. The root was set to the SOX2 expression using scf.tl.root and corresponded to the pMN. Later, the pseudotime was computed using scf.tl.pseudotime with n_jobs=20, n_map=50, seed=42. To explore the differences in motoneuron differentiation along a shared trajectory, the data were stratified by genotype into CTRL and SMA MN groups. For each group, dendrograms were generated using scf.pl.dendrogram and KDE plots (seaborn.kdeplot). Next, the pySCENIC AUCell output was processed to align with the motoneuron trajectory data generated before and this approach was adopted from the study of Petitpré C et al.2022^74^. pySCENIC AUCell output for control and SMA samples was loaded from .loom files. The AUCell matrices were aligned with the trajectory data by matching cell IDs. Regulons active in fewer than six cells were filtered out. Regulon activity was tested for association with trajectory data using scFates and GAM (scf.tl.test_association with n_jobs=20 and A_cut=0.025). Pseudotime for cells was updated based on their order in trajectory analysis (adata.uns["pseudotime_list"]), and the trajectory model was fitted (scf.tl.fit with n_jobs=20). The trend of specific regulons activity along the trajectory segments was visualised using scf.pl.single_trend and scf.pl.trends. For control samples, a set of significant regulons was highlighted, and the trends of these regulons were plotted. Similar analyses were performed for SMA samples.

#### Identification of informative splicing gene modules

To associate splicing gene modules to clusters in the SCOs object (CTRL or SMA), we subsetted the object to contain only the selected pools of splicing genes but all cells. Using scVI dimensionality reduction, Hotspot^74^ (v=1.1.1) analysis was conducted to identify gene modules by calculating autocorrelations for the genes and identifying significant genes with an FDR < 0.05. Local correlations were computed for the significant genes using 15 parallel jobs, and gene modules were created with a minimum gene threshold of 50, core-only mode, and an FDR threshold of 0.05. Module scores were calculated and assigned to the dataset. Finally, module scores of splicing genes were visualised using scatter plots on UMAP coordinates with custom color palettes.

#### Correlation to fetal spinal cord data

Single-cell RNA sequencing datasets from SCOs object and first and second trimester fetal spinal cord tissue^23^ were loaded as AnnData objects. The organoid dataset was subset to include only control samples (“CTRL”) for subsequent correlation analysis. Unique identifiers for each observation were ensured using the obs_names_make_unique() function. To minimize noise, genes were filtered by selecting the top 2000 highly variable genes (HVGs) for each dataset using Scanpy’s highly_variable_genes() with the seurat_v3 method. The datasets were normalized to a target count of 10,000 per cell and log-transformed using sc.pp.normalize_total() and sc.pp.log1p() respectively. The first and second trimester fetal datasets were labeled by gestational week (GW) and grouped by regions for analysis. Mean expression values per group were calculated using a custom function to generate a grouped mean expression matrix for each dataset. Common genes between the fetal and organoid datasets were identified, and Pearson correlations were calculated between fetal regions and organoid CTRL genotype to quantify similarity in expression profiles. Two correlation matrices, one for each trimester, were computed and saved for later analysis. The correlation matrices were subsequently combined, and a heatmap of the Pearson correlation coefficients was visualized using Seaborn. The combined correlation heatmap was used to interpret relationships between fetal regions and the control organoid samples, focusing on similarities and differences across developmental stages and phenotypic profiles.

### Bulk RNA-seq analyses

#### Library construction

Libraries were prepared starting from 50 ng of total RNA derived from 9 control, 6 SMA, and 16 morpholino-treated SMA (8 r6-MO, 8 r6-MO-scrambled) organoids, using the SMARTer Stranded RNA-Seq Kit, TaKaRa) and following the manufacturer’s instructions. Samples underwent a high coverage paired-end 150-bp strand-specific sequencing using a NextSeq 2000 platform (Illumina).

#### Alignment

For each sample (9x control organoids, 6x SMA organoids, 8x SMA organoids treated with r6-MO and 8x SMA organoids treated with r6-MO-Scramble), raw reads were mapped to the human hg38 genome and counted using STAR v2.7.10a^75^.

#### *SMN1* and *SMN2* counts

*SMN1* (SMN-C) and *SMN2* (SMN-T) genes are characterized by a very high degree of homology (more than 99%). Thus, to distinguish the reads derived from the two genes we counted the number of reads mapping to exon 7, where the presence of a nucleotide difference between the two genes enables their unambiguous identification. We performed this check using the IGV tool. Counts mapping elsewhere on the *SMN* genes were not considered for this analysis.

#### Differential gene expression analysis

Differential gene expression analysis was carried out using R and the edgeR package v4.0.2 with the functions estimateDisp, glmFit and glmLRT. Genes having cpm < 0.05 in at least 6 samples were filtered out.

#### Gene ontology Differential Expressed Genes

Gene ontology of differential expressed genes between CTRL and SMA organoids was done using Ingenuity Pathway Analysis. Terms from “Canonical pathways” and “diseases and functions” were filtered.

#### Differential splicing analysis

Differential splicing analysis between samples was determined using rMATS turbo v4.3.0^76^, a tool that identifies 5 basic types of AS patterns: skipped exons (SE), mutually exclusive exons (MXE), alternative 5’ (A5SS) and 3’ splice sites (A3SS), and retained introns (RI). Significant events were characterized by a False Discovery Rate (FDR) < 0.05 and an absolute Inclusion Level Difference > 0.05. Only the reads mapping to exon-exon junctions were used for the analyses. Sashimi plot of events of interest were created using rmats2sashimiplot v2.0.4 (https://github.com/Xinglab/rmats2sashimiplot).

#### Gene ontology of splicing events

Differential splicing events from SMA vs CTRL and SMA r6-MO vs SMA r6-MO-Scrambled were compared. Events sharing (i) the same starting position and the same ending position and (ii) with opposite Inclusion Level Difference were called as “common events”. To annotate gene function, gene ontology was performed using Metascape^77^ (^5^). Downstream processing of splicing and gene expression data were done in R v4.1.0 using custom scripts.

### Statistical analysis

Continuous variables were reported as mean±SD. Comparisons between groups and between baseline and post-intervention or interval within a group were assessed using unpaired and paired two-sided Student’s *t-test*, respectively. One-way or two-way ANOVA was used for comparisons among groups, with Tukey HSD or Sidak’s post-hoc tests applied when the null hypothesis for the omnibus analysis was rejected. GraphPad Prism v.8 was used for all analyses with a significance level of 0.05.

## Acknowledgments

The authors thank Associazione Centro Dino Ferrari for support, and the Humanitas Genomic Unit (HUGE) and Humanitas Bioinformatic Unit for technical support in single-cell sequencing experiments and analysis. We acknowledge Teresa Rayon (Babraham Institute, Cambridge, UK) for sharing counts of human fetal pMN and MN cell populations and Hynek Wichterle for sharing HB9 antibody (Motor Neuron Center, Columbia university, New York, NY). We acknowledge Dario Ronchi for precious comments and Stefania Sempreboni for technical support.

## Authors’ contributions

S.C., M.N., S.L, I.F., M.T. and P.R. conceived and designed the research project. M.N, P.R., M.R., G.F., A.D.A. and L.C. performed experiments and collected data regarding iPSCs generation and characterization. P.R, I.F. and A.D.A. generated and characterized spinal cord organoids. F.B. helped generating and characterizing spinal cord organoids. M.T, P.R and I.F generated and characterized brain organoids. S.M. and M.T. performed image acquisition and S.M., M.T. and P.R. performed image quantification. L.M. and E.D. conducted MEA recording and analyses with the help of E.P., and C.C. performed and analyzed calcium imaging data and TUNEL quantification together with P.R. I.F. and P.R. performed statistical analyses on aggregated data. C.P., S.M., and M.T. performed single-cell RNA-seq experiments and P.K. and I.S. performed the relative computational analyses. M. M. and E.M.P. analyzed bulk RNA-seq and splicing data. G.C. contributed to data discussion and supervision of I.F. and P.R. I.F., P.R., M.T., S.L., and S.C. wrote the manuscript with the critical contribution of all the authors. All authors critically reviewed and approved the final version of the manuscript.

## Supplementary material

**Supplementary Figure 1.** SCO molecular profiling.

**Supplementary Figure 2.** CTRL and SMA SCO molecular profiling and differential analyses.

**Supplementary Figure 3.** CTRL and SMA SCO apoptosis analyses.

**Supplementary Figure 4.** CTRL and SMA SCO MN partition.

**Supplementary Figure 5.** CTRL and SMA SCOs present active mechanisms of neuronal inhibition.

**Supplementary Figure 6.** CTRL and SMA cerebral organoid characterization.

**Supplementary Figure 7.** Gene expression and alternative splicing events in CTRL, SMA and r6-MO treated SMA SCOs.

References ^78,79^.

## Supplementary Figures

**Supplementary Figure 1. CTRL and SMA molecular profiling and differential analyses.**

**A**, UMAP representation of the aggregate data from 2 months old SCOs, four CTRL SCOs (n=2 lines from C3 and n=2 lines from C6) and four SMA SCOs (n=2 lines from S3 and n=2 lines from S4). **B**, Frequency of cells per cluster for CTRL and SMA SCOs. **C**, Heatmap illustrating the Pearson correlation coefficients between aggregated control samples and fetal spinal cord samples from the first and second trimesters, categorized by region. **D,** Dot plots illustrating the expression of representative genes associated with anterior and posterior genes in different cell types. **E**, Violin plots showing module score enrichment in individual clusters for selected relevant cell populations. UMAP feature plots showing the log-normalized counts of selected gene modules for cycling cells, progenitors, floor plate progenitor (pFP), unmatched-SMA progenitors (UnSMA-P), astroglia, neurons, V2a and V2b neurons.

**Supplementary Figure 2. Differential gene expression analysis between SMA and CTRL in other identified clusters.**

**A**, Volcano plot of differential gene expression in individual clusters in SMA SCOs relative to CTRL in astroglia, pFP, pMN, V2a and V2b. Values of log2 fold change are plotted against - log10 of the q-value for each gene. Red denotes genes upregulated in SMA while blue denotes genes downregulated in SMA (absolute log fold change ≥ 0.2 min.pct = 0.25 and adjusted *q*- value < 0.05, thresholds plotted as dashed lines). The bar plot illustrates the selected Gene Ontology (GO) terms associated with disease and canonical pathways, derived from both upregulated and downregulated genes. These GO terms were selected based on their relevance to the observed gene expression changes, when present, in the corresponding cluster performed with g:Profiler (bar plots with color of the associated false discovery rate (FDR)).

**Supplementary Figure 3. CTRL and SMA SCO differentiation trajectories.**

**A**, Representative images of TUNEL (green) staining of CTRL and SMA SCOs at 1 and 2 mo. Scale bar: 100 µm. Nuclei were stained with DAPI (blue). **B**, Quantification of TUNEL-positive cells (as a percentage of cells revealed by DAPI staining) at 1 (circles) and 2 mo (triangles) (two-way ANOVA, Sidak’s multiple comparisons test: *p*= 0.3786 and *p*< 0.0001 for CTRL and SMA, respectively). The proportion of TUNEL positive cells increased significantly at 2 mo in comparison with 1 mo in SMA but not CTRL SCOs. **C**, Dot plot showing apoptosis-related biological function enrichment analysis based on differentially expressed genes (pseudobulk DGE) between SMA and CTRL SCOs for different cell types (columns). Dot color represents z-score (blue: negative, red: positive), and dot size reflect the significance.

**Supplementary Figure 4. Differential gene expression analysis between SMA and CTRL in MN clusters.**

**A**-**B,** UMAP plot of the subclustering of immature and mature MNs in CTRL and SMA SCOs. **C**, UMAP feature plots showing the log-normalized counts of selected representative genes for neuronal progenitors (*SOX2*, *NKX6-1* and *OLIG2*), and neuronal differentiation markers (*ASCL1, NEUROG2, NEUROG4, PHOX2B, PHOX2A, PNT, MNX1, ISL1* and *ELAVL2*).

**Supplementary Figure 5. CTRL and SMA SCOs present active mechanisms of neuronal inhibition.**

**A**, Heatmap showing the average expression levels of glutamate receptor genes within CTRL and SMA SCOs in neuronal clusters. **B**, Representative images of GABA (upper panel, red) and GAD65 (lower panel, red) staining in CTRL and SMA SCOs. Scale bar: upper panel 100 µm, lower panel 50 µm. Nuclei were stained with DAPI (blue). **C**, Quantification of the change in firing frequency (%) in reactive channels after gabazine perfusion (unpaired t-test, *p*=0.9521). **D**, Quantification of reactive channels (%) after gabazine and strychnine perfusion (unpaired t-test, *p*=0.5455).

**Supplementary Figure 6. Characterization and functional analysis of CTRL and SMA CrOs.**

**A**, Staining of CTRL (left panel) and SMA (right panel) CrOs at 2 mo. Nuclei were stained with DAPI (blue). Scale bar: 50 µm. **B**, Whole mount staining of CTRL and SMA CrOs revealed elongated filaments of GFAP-positive cells and the post-mitotic neuronal marker TUJ1. Scale bar: 50 µm. **C**, Representative image of CTRL CrOs under different conditions after Fluo4 loading. Calcium imaging showed spatially organized enhanced calcium flux after glutamate (glu) administration, which increased upon ionomycin (iono) administration and decreased after EDTA chelation, which was used as controls for intracellular calcium visualization. Scale bar: 500 µm. **D**, Representative images of GAD65 staining in CTRL and SMA CrOs. Scale bar: 20 µm. **E**, Upper panel: Quantification of reactive channels (%) after gabazine and strychnine perfusion (unpaired t-test, *p*=0.9521). Lower panel: Quantification of the change in firing frequency (%) in reactive channels after gabazine perfusion (unpaired t-test, *p*=0.1452). **F**, Representative image of CTRL and SMA cerebral organoids after Fluo4 loading in basal condition (left) and after adding glutamate (right). **G**, Maximum amplitude of calcium variation in CTRL and SMA organoids in basal conditions *vs.* glutamate-stimulated conditions (paired t-tests: *p*=0.0315 and *p*=0.0313 for CTRL and SMA respectively). **H**, Relative change in maximum amplitude variation in basal *vs.* stimulated conditions in organoids from (**G**) (unpaired t-test, *p*=0.02325).

**Supplementary Figure 7. RNA-sequencing identifies gene expression and alternative splicing events changes between CTRL, SMA and r6-MO treated SMA SCOs.**

**A**, Volcano plot of differential gene expression changes between SMA and CTRL SCOs. **B**, Selected Gene Ontology terms of the 2477 differentially expressed genes between SMA and CTRL SCOs showing the -log10(p-value) and colored by z-score. Analysis was performed with QIAGEN Ingenuity Pathway Analysis (IPA) software. **C**, Clustermap showing the single-cell co-expression profiles of genes with altered expression in SMA *vs.* CTRL SCOs organized into distinct modules. **D**, Summary of the total number of splicing events measured between SMA/CTRL (grey) and r6-MO SMA/r6-Scramble SMA (yellow) and the total number of associated genes per splicing event category. **E**, Volcano plot of differentially expressed genes between unmatched SMA progenitors and CTRL progenitors in SCOs. **F**, Volcano plot of differentially expressed genes between unmatched SMA neurons and CTRL neurons in SCOs. In both plots (**E**, **F**), points represent genes, with the x-axis showing log2 fold change and the y-axis showing –log10(p-value). Gene names of significantly reverted genes identified in the splicing analysis are highlighted.

## References

1. Kolb, S. J. et al. Natural history of infantile-onset spinal muscular atrophy. Ann Neurol 82, 883–891 (2017).

2. Faravelli, I., Nizzardo, M., Comi, G. P. & Corti, S. Spinal muscular atrophy--recent therapeutic advances for an old challenge. Nat Rev Neurol 11, 351–359 (2015).

3. Lorson, C. L., Hahnen, E., Androphy, E. J. & Wirth, B. A single nucleotide in the SMN gene regulates splicing and is responsible for spinal muscular atrophy. Proceedings of the National Academy of Sciences 96, 6307–6311 (1999).

4. Parente, V. & Corti, S. Advances in spinal muscular atrophy therapeutics. Ther Adv Neurol Disord 11, 1756285618754501 (2018).

5. Mercuri, E., Pera, M. C., Scoto, M., Finkel, R. & Muntoni, F. Spinal muscular atrophy - insights and challenges in the treatment era. Nat Rev Neurol 16, 706–715 (2020).

6. Mercuri, E., Sumner, C. J., Muntoni, F., Darras, B. T. & Finkel, R. S. Spinal muscular atrophy. Nat Rev Dis Primers 8, 52 (2022).

7. Soler-Botija, C., Ferrer, I., Gich, I., Baiget, M. & Tizzano, E. F. Neuronal death is enhanced and begins during foetal development in type I spinal muscular atrophy spinal cord. Brain 125, 1624–1634 (2002).

8. Ramos, D. M. et al. Age-dependent SMN expression in disease-relevant tissue and implications for SMA treatment. J Clin Invest 129, 4817–4831 (2019).

9. Kong, L. et al. Impaired prenatal motor axon development necessitates early therapeutic intervention in severe SMA. Sci Transl Med 13, eabb6871 (2021).

10. Mendonça, R. H. et al. Severe brain involvement in 5q spinal muscular atrophy type 0. Ann Neurol 86, 458–462 (2019).

11. Faravelli, I. & Corti, S. Spinal muscular atrophy - challenges in the therapeutic era. Nat Rev Neurol 16, 655–656 (2020).

12. Tiziano, F. D. & Tizzano, E. F. 25 years of the SMN genes: the Copernican revolution of spinal muscular atrophy. Acta Myol 39, 336–344 (2020).

13. Tizzano, E. F. et al. Outcomes for patients in the RESTORE registry with spinal muscular atrophy and four or more SMN2 gene copies treated with onasemnogene abeparvovec. Eur J Paediatr Neurol 53, 18–24 (2024).

14. Lancaster, M. A. et al. Cerebral organoids model human brain development and microcephaly. Nature 501, 373–379 (2013).

15. Velasco, S. et al. Individual brain organoids reproducibly form cell diversity of the human cerebral cortex. Nature 570, 523–527 (2019).

16. Andersen, J. et al. Generation of Functional Human 3D Cortico-Motor Assembloids. Cell 183, 1913–1929.e26 (2020).

17. Ma, X. R. et al. TDP-43 represses cryptic exon inclusion in the FTD–ALS gene UNC13A. Nature 603, 124–130 (2022).

18. Sagner, A. & Briscoe, J. Establishing neuronal diversity in the spinal cord: a time and a place. Development 146, dev182154 (2019).

19. Lupo, G., Harris, W. A. & Lewis, K. E. Mechanisms of ventral patterning in the vertebrate nervous system. Nat Rev Neurosci 7, 103–114 (2006).

20. Amoroso, M. W. et al. Accelerated high-yield generation of limb-innervating motor neurons from human stem cells. J Neurosci 33, 574–586 (2013).

21. Du, Z.-W. et al. Generation and expansion of highly pure motor neuron progenitors from human pluripotent stem cells. Nat Commun 6, 6626 (2015).

22. Rizzo, F. et al. Key role of SMN/SYNCRIP and RNA-Motif 7 in spinal muscular atrophy: RNA-Seq and motif analysis of human motor neurons. Brain 142, 276–294 (2019).

23. Zhang, Q. et al. Single-cell analysis reveals dynamic changes of neural cells in developing human spinal cord. EMBO reports 22, e52728 (2021).

24. Dann, E., Henderson, N. C., Teichmann, S. A., Morgan, M. D. & Marioni, J. C. Differential abundance testing on single-cell data using k-nearest neighbor graphs. Nat Biotechnol 40, 245–253 (2022).

25. Faure, A. J. et al. Mapping the energetic and allosteric landscapes of protein binding domains. Nature 604, 175–183 (2022).

26. Rayon, T., Maizels, R. J., Barrington, C. & Briscoe, J. Single-cell transcriptome profiling of the human developing spinal cord reveals a conserved genetic programme with human-specific features. Development 148, dev199711 (2021).

27. Sagner, A. et al. A shared transcriptional code orchestrates temporal patterning of the central nervous system. PLoS Biol 19, e3001450 (2021).

28. Van de Sande, B. et al. A scalable SCENIC workflow for single-cell gene regulatory network analysis. Nat Protoc 15, 2247–2276 (2020).

29. Meng, H. et al. Loss of Parkinson’s disease-associated protein CHCHD2 affects mitochondrial crista structure and destabilizes cytochrome c. Nat Commun 8, 15500 (2017).

30. Kee, T. R. et al. Mitochondrial CHCHD2: Disease-Associated Mutations, Physiological Functions, and Current Animal Models. Front Aging Neurosci 13, 660843 (2021).

31. Arumugam, S., Garcera, A., Soler, R. M. & Tabares, L. Smn-Deficiency Increases the Intrinsic Excitability of Motoneurons. Front Cell Neurosci 11, 269 (2017).

32. Quinlan, K. A. et al. Hyperexcitability precedes motoneuron loss in the Smn2B/- mouse model of spinal muscular atrophy. J Neurophysiol 122, 1297–1311 (2019).

33. Brunet, A., Stuart-Lopez, G., Burg, T., Scekic-Zahirovic, J. & Rouaux, C. Cortical Circuit Dysfunction as a Potential Driver of Amyotrophic Lateral Sclerosis. Front Neurosci 14, 363 (2020).

34. Sibilla, S. & Ballerini, L. GABAergic and glycinergic interneuron expression during spinal cord development: dynamic interplay between inhibition and excitation in the control of ventral network outputs. Prog Neurobiol 89, 46–60 (2009).

35. Ramirez, A. et al. Investigation of New Morpholino Oligomers to Increase Survival Motor Neuron Protein Levels in Spinal Muscular Atrophy. Int J Mol Sci 19, 167 (2018).

36. Moulton, H. M. & Moulton, J. D. Morpholinos and their peptide conjugates: therapeutic promise and challenge for Duchenne muscular dystrophy. Biochim Biophys Acta 1798, 2296–2303 (2010).

37. Bersani, M. et al. Cell-penetrating peptide-conjugated Morpholino rescues SMA in a symptomatic preclinical model. Mol Ther 30, 1288–1299 (2022).

38. Gubitz, A. K., Feng, W. & Dreyfuss, G. The SMN complex. Exp Cell Res 296, 51–56 (2004).

39. Yoo, H. et al. ATP-Dependent Chromatin Remodeler CHD9 Controls the Proliferation of Embryonic Stem Cells in a Cell Culture Condition-Dependent Manner. Biology 9, 428 (2020).

40. Guo, W. et al. HDAC6 inhibition reverses axonal transport defects in motor neurons derived from FUS-ALS patients. Nat Commun 8, 861 (2017).

41. Chen, S. et al. Histone deacetylase 6 delays motor neuron degeneration by ameliorating the autophagic flux defect in a transgenic mouse model of amyotrophic lateral sclerosis. Neurosci. Bull. 31, 459–468 (2015).

42. Mancinelli, S. & Lodato, S. Decoding neuronal diversity in the developing cerebral cortex: from single cells to functional networks. Curr Opin Neurobiol 53, 146–155 (2018).

43. Velasco, S., Paulsen, B. & Arlotta, P. 3D Brain Organoids: Studying Brain Development and Disease Outside the Embryo. Annu Rev Neurosci 43, 375–389 (2020).

44. Tambalo, M. & Lodato, S. Brain organoids: Human 3D models to investigate neuronal circuits assembly, function and dysfunction. Brain Res 1746, 147028 (2020).

45. Paulsen, B. et al. Autism genes converge on asynchronous development of shared neuron classes. Nature 602, 268–273 (2022).

46. Antón-Bolaños, N. et al. Brain Chimeroids reveal individual susceptibility to neurotoxic triggers. Nature 631, 142–149 (2024).

47. Ogura, T., Sakaguchi, H., Miyamoto, S. & Takahashi, J. Three-dimensional induction of dorsal, intermediate and ventral spinal cord tissues from human pluripotent stem cells. Development 145, dev162214 (2018).

48. Faustino Martins, J.-M., et al. Self-Organizing 3D Human Trunk Neuromuscular Organoids. Cell Stem Cell 26, 172–186.e6 (2020).

49. Galimberti, M. et al. Huntington’s disease cellular phenotypes are rescued non-cell autonomously by healthy cells in mosaic telencephalic organoids. Nat Commun 15, 6534 (2024).

50. Barnat, M. et al. Huntington’s disease alters human neurodevelopment. Science 369, 787–793 (2020).

51. Galgoczi, S. et al. Huntingtin CAG expansion impairs germ layer patterning in synthetic human 2D gastruloids through polarity defects. Development 148, dev199513 (2021).

52. Motyl, A. A. L. et al. Pre-natal manifestation of systemic developmental abnormalities in spinal muscular atrophy. Hum Mol Genet 29, 2674–2683 (2020).

53. Tharaneetharan, A. et al. Functional Abnormalities of Cerebellum and Motor Cortex in Spinal Muscular Atrophy Mice. Neuroscience 452, 78–97 (2021).

54. Simon, C. M. et al. A Stem Cell Model of the Motor Circuit Uncouples Motor Neuron Death from Hyperexcitability Induced by SMN Deficiency. Cell Rep 16, 1416–1430 (2016).

55. Wainger, B. J. et al. Intrinsic membrane hyperexcitability of amyotrophic lateral sclerosis patient-derived motor neurons. Cell Rep 7, 1–11 (2014).

56. Burg, T. et al. Absence of Subcerebral Projection Neurons Is Beneficial in a Mouse Model of Amyotrophic Lateral Sclerosis. Ann Neurol 88, 688–702 (2020).

57. Marques, C., Burg, T., Scekic-Zahirovic, J., Fischer, M. & Rouaux, C. Upper and Lower Motor Neuron Degenerations Are Somatotopically Related and Temporally Ordered in the Sod1 Mouse Model of Amyotrophic Lateral Sclerosis. Brain Sci 11, 369 (2021).

58. Finkel, R. S. et al. Nusinersen versus Sham Control in Infantile-Onset Spinal Muscular Atrophy. New England Journal of Medicine 377, 1723–1732 (2017).

59. Duan, D., Goemans, N., Takeda, S., Mercuri, E. & Aartsma-Rus, A. Duchenne muscular dystrophy. Nat Rev Dis Primers 7, 1–19 (2021).

60. Corti, S. et al. Genetic correction of human induced pluripotent stem cells from patients with spinal muscular atrophy. Sci Transl Med 4, 165ra162 (2012).

61. Carpenter, A. E. et al. CellProfiler: image analysis software for identifying and quantifying cell phenotypes. Genome Biol 7, R100 (2006).

62. Schindelin, J., et al. Fiji: an open-source platform for biological-image analysis. Nat Methods 9, 676–682 (2012).

63. Monteverdi, A., Di Domenico, D., D’Angelo, E. & Mapelli, L. Anisotropy and Frequency Dependence of Signal Propagation in the Cerebellar Circuit Revealed by High-Density Multielectrode Array Recordings. Biomedicines 11, 1475 (2023).

64. Wolf, F. A., Angerer, P. & Theis, F. J. SCANPY: large-scale single-cell gene expression data analysis. Genome Biology 19, 15 (2018).

65. Wolock, S. L., Lopez, R. & Klein, A. M. Scrublet: Computational Identification of Cell Doublets in Single-Cell Transcriptomic Data. Cell Syst 8, 281–291.e9 (2019).

66. Gayoso, A. et al. A Python library for probabilistic analysis of single-cell omics data. Nat Biotechnol 40, 163–166 (2022).

67. Traag, V. A., Waltman, L. & van Eck, N. J. From Louvain to Leiden: guaranteeing well-connected communities. Sci Rep 9, 5233 (2019).

68. Ahlmann-Eltze, C. & Huber, W. Comparison of transformations for single-cell RNA-seq data. Nat Methods 20, 665–672 (2023).

69. Muzellec, B., Teleńczuk, M., Cabeli, V. & Andreux, M. PyDESeq2: a python package for bulk RNA-seq differential expression analysis. Bioinformatics 39, btad547 (2023).

70. Badia-i-Mompel, P. et al. decoupleR: ensemble of computational methods to infer biological activities from omics data. Bioinformatics Advances 2, vbac016 (2022).

71. Setty, M. et al. Characterization of cell fate probabilities in single-cell data with Palantir. Nat Biotechnol 37, 451–460 (2019).

72. Moerman, T. et al. GRNBoost2 and Arboreto: efficient and scalable inference of gene regulatory networks. Bioinformatics 35, 2159–2161 (2019).

73. AUCell. Bioconductor http://bioconductor.org/packages/AUCell/.

74. Petitpré, C. et al. Single-cell RNA-sequencing analysis of the developing mouse inner ear identifies molecular logic of auditory neuron diversification. Nat Commun 13, 3878 (2022).

75. Dobin, A. et al. STAR: ultrafast universal RNA-seq aligner. Bioinformatics 29, 15–21 (2013).

76. Shen, S. et al. rMATS: robust and flexible detection of differential alternative splicing from replicate RNA-Seq data. Proc Natl Acad Sci U S A 111, E5593–5601 (2014).

77. Zhou, Y. et al. Metascape provides a biologist-oriented resource for the analysis of systems-level datasets. Nat Commun 10, 1523 (2019).

78. Yin, X., Xia, J., Sun, Y. & Zhang, Z. CHCHD2 is a potential prognostic factor for NSCLC and is associated with HIF-1a expression. BMC Pulmonary Medicine 20, 40 (2020).

79. Sumner, C. J. et al. SMN mRNA and protein levels in peripheral blood: biomarkers for SMA clinical trials. Neurology 66, 1067–1073 (2006).

