## Supplementary Figures for "Targeted Antisense Oligonucleotide Treatment Rescues Developmental Alterations in Spinal Muscular Atrophy Organoids"

### Supplementary Figure 1

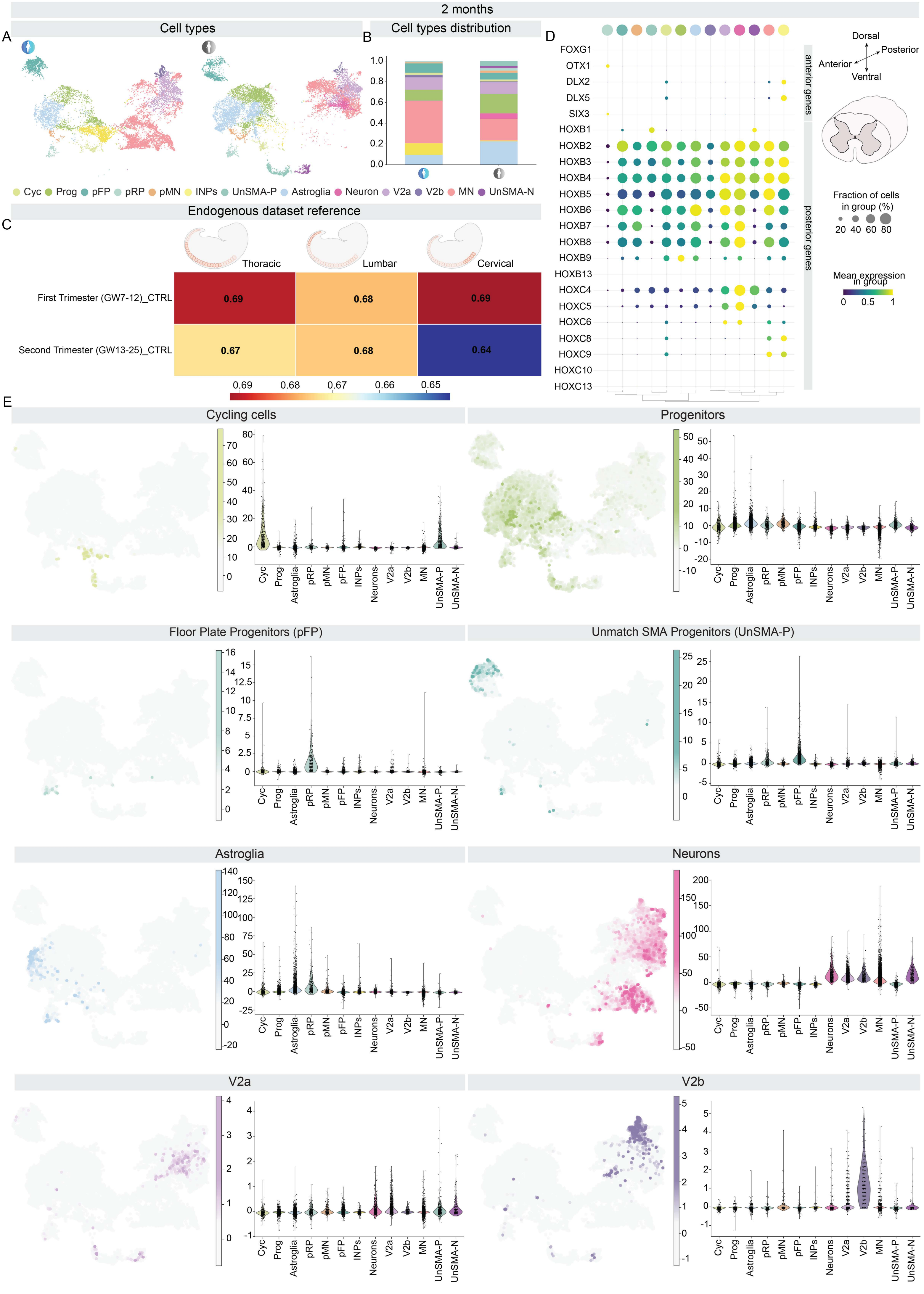

#### Supplementary Figure 2

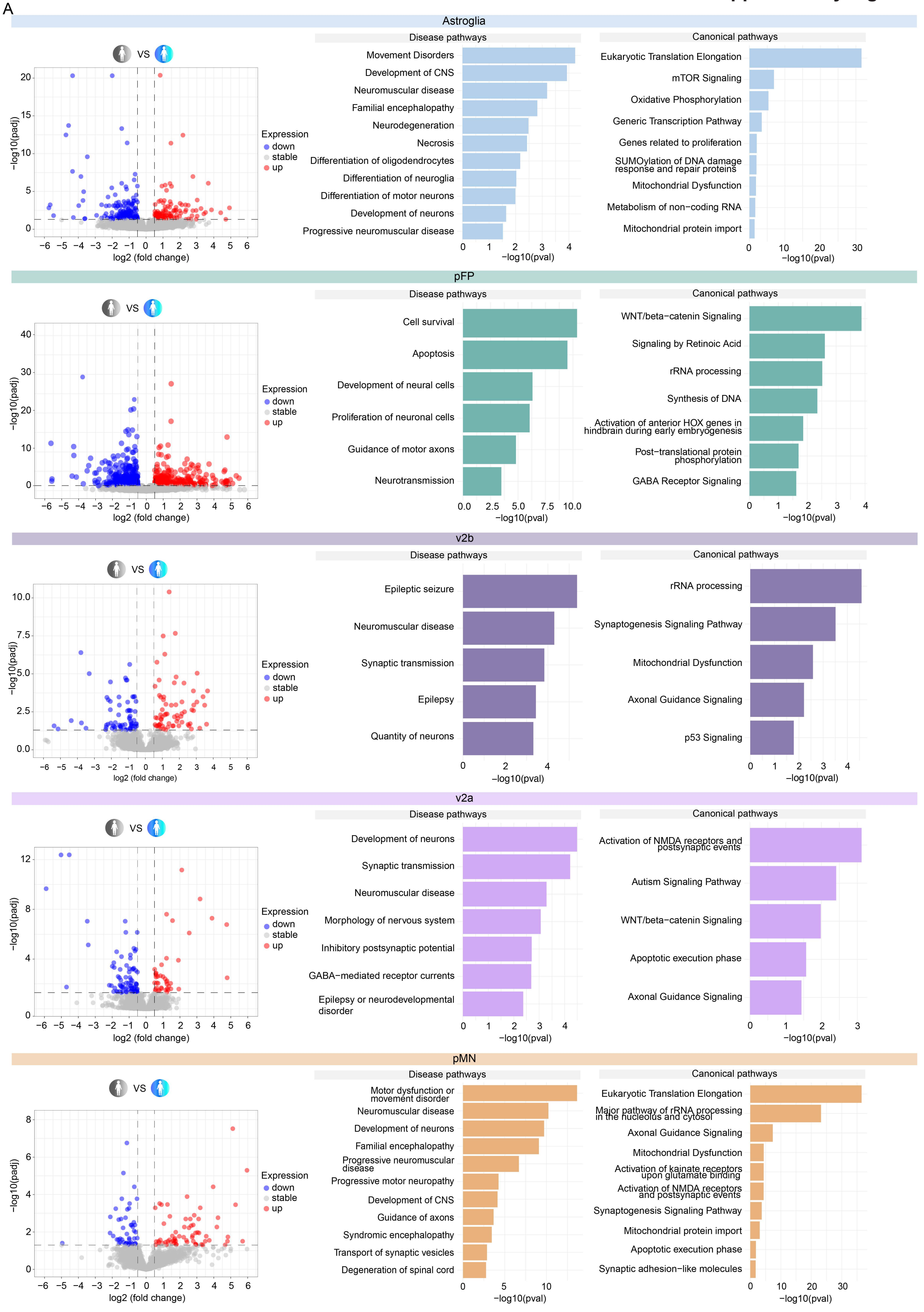

### Supplementary Figure 3

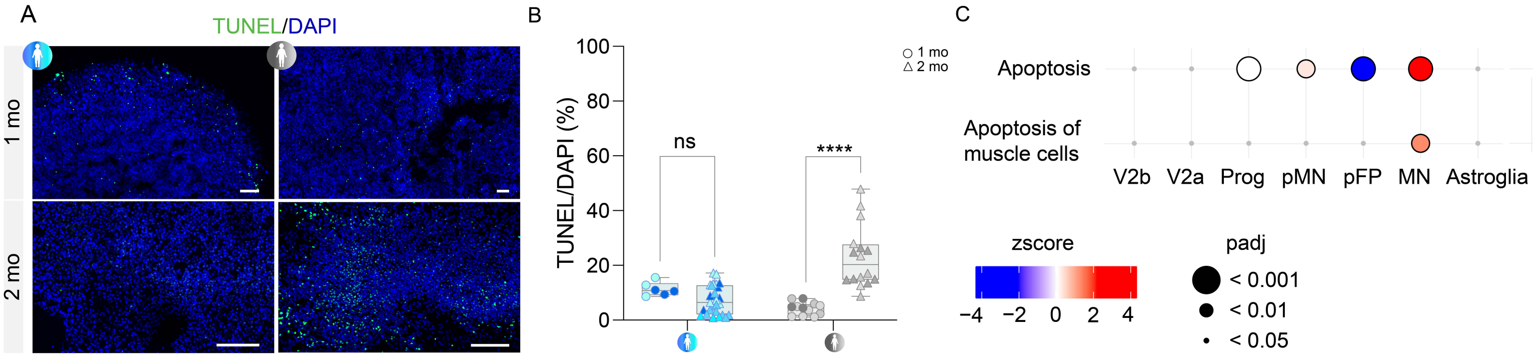

#### Supplementary Figure 4

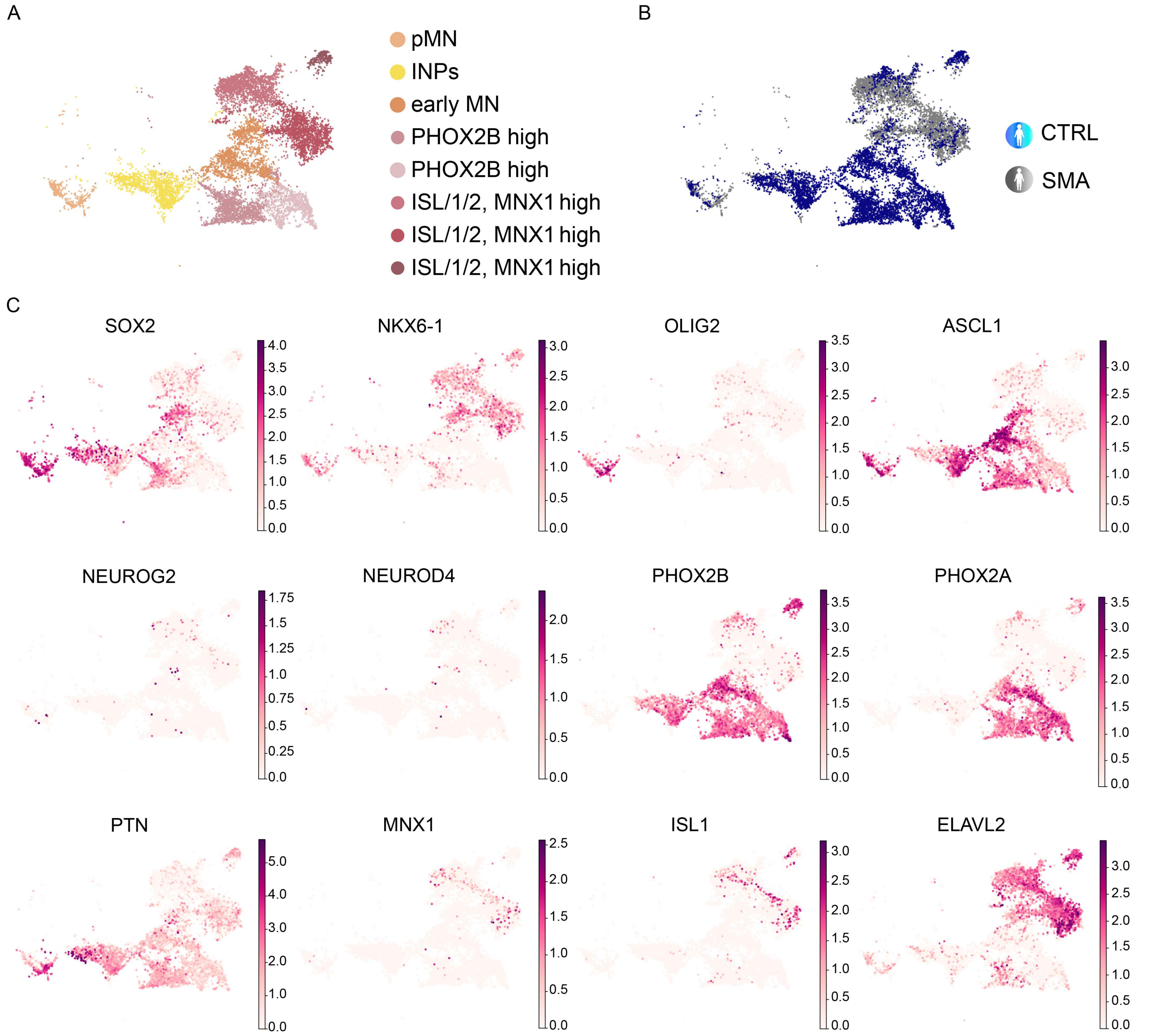

### Supplementary Figure 5

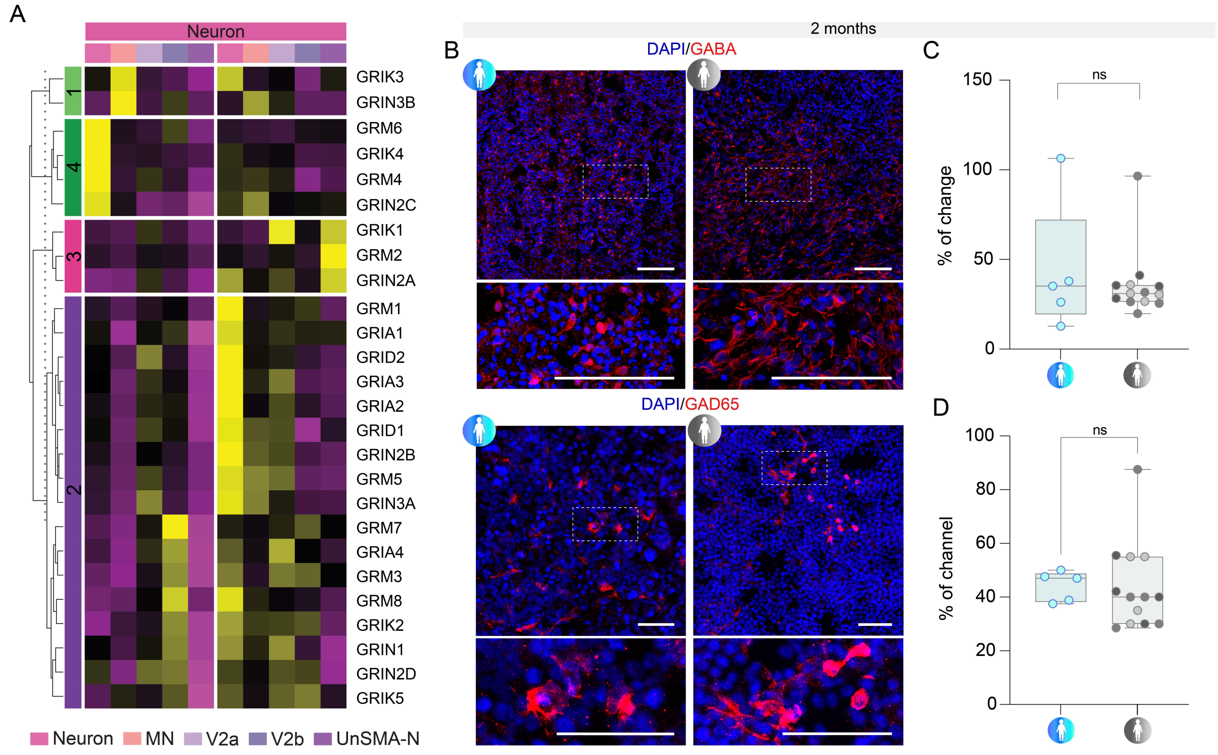

### Supplementary Figure 6

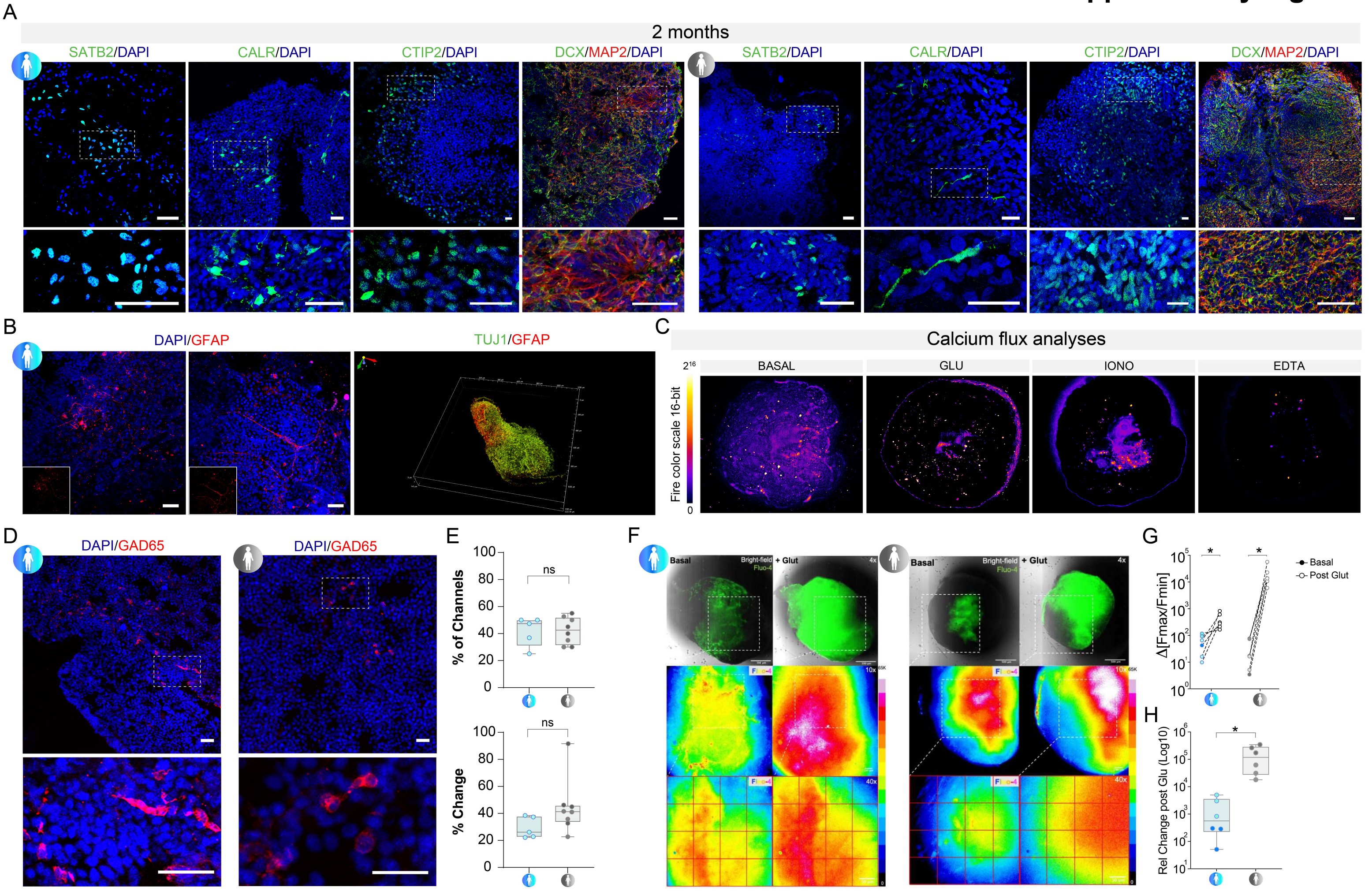

### Supplementary Figure 7

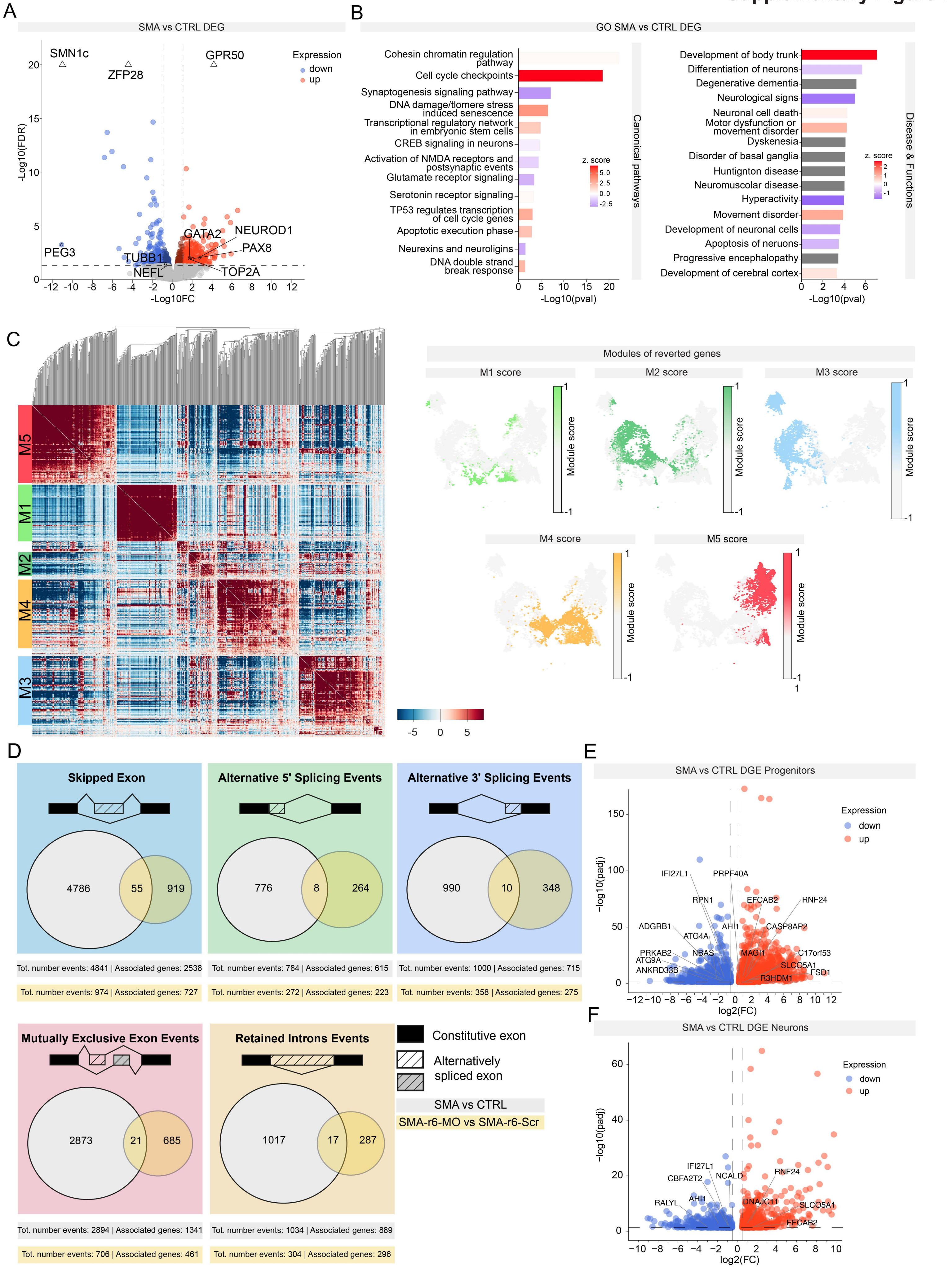
